# Alcohol dehydrogenase-mediated methanol dissimilation increases carbon efficiency in synthetic autotrophic yeast

**DOI:** 10.64898/2026.03.09.710585

**Authors:** Charles Moritz, Lisa Lutz, Michael Baumschabl, David Glinsner, Thomas Gassler, Diethard Mattanovich, Özge Ata

## Abstract

The efficient production of food and biochemicals using microorganisms that utilize single-carbon feedstocks presents a promising approach for advancing a circular bioeconomy. *Komagataella phaffii* (formerly *Pichia pastoris*) is a methylotrophic yeast already widely used in industry, making it an attractive host for such applications. Recently, *K. phaffii* was converted into an autotrophic strain capable of assimilating CO_2_ into both biomass and secreted organic acids, using energy derived from dissimilation of methanol to CO_2_. In these strains, methanol oxidation is catalysed by an alcohol oxidase (Aox2), which transfers electrons to oxygen without conserving reducing equivalents. To address this limitation, in this study we explored redirecting methanol dissimilation through the native alcohol dehydrogenase (Adh2), coupling methanol oxidation with NADH generation to improve carbon efficiency. By deleting *AOX2* and overexpressing *ADH2*, we generated Adh2-based autotrophic strains that exhibited growth rates comparable to the parental strain (0.007 h⁻¹), while reducing specific CO₂ production by 53% and increasing biomass yield (Y_X/MeOH_) by 59%. We further applied this strategy to convert previously developed autotrophic strains producing itaconic acid and lactic acid into Adh2-dependent strains. Optimizing *ADH2* expression through multicopy integration resulted in strains with approximately two-fold higher molar carbon efficiency (Y_(X+P)/CO2_) while achieving elevated product titers—2.2-fold for itaconic acid and 3.8-fold for lactic acid—relative to the parental strains. Our findings demonstrate that alcohol dehydrogenase-mediated methanol dissimilation can significantly improve yield and productivity of autotrophic *K. phaffii* strains, with broad implications for sustainable bioproduction from one-carbon substrates.

## Introduction

Anthropogenic greenhouse gas emissions and land-use change are driving global climate warming, posing major challenges to both human societies and natural ecosystems^1^. Since the pre-industrial era, atmospheric CO₂ concentrations have increased by over 50%, reaching 419 ppm in 2023, while annual global greenhouse gas emissions continue to rise^2^. Global surface temperatures have continued to rise at an accelerating pace, with estimates for 2024 exceeding the 1.5°C threshold set by the 2016 Paris Agreement^3^. At the same time, demand for materials such as plastic polymers remains high and is currently met predominantly by fossil fuel-derived feedstocks. The use of captured CO₂ as a substrate for microbial conversion to biopolymers offers a strategy to both mitigate CO₂ emissions and reduce reliance on petrochemical resources, without competing for arable land^4^. As CO_2_ is a highly oxidized substrate, autotrophic organisms require an external source of reducing power. In microalgae and plants, this reducing power is provided as NADPH through light-dependent photosynthesis. Many microorganisms have evolved alternative metabolic strategies that fix CO₂ by harvesting reducing equivalents from high-energy, reduced compounds such as hydrogen, formate, and methanol, thereby driving CO₂ fixation while simultaneously supplying cellular energy^5,6^. When produced using renewable energy, these co-substrates provide a promising basis for the emerging single-carbon (C1) bioeconomy^7,8,9^.

Recent advances in metabolic engineering allow metabolic traits to be transferred between microorganisms via heterologous pathway expression. In synthetic autotrophy, this strategy has enabled the conversion of heterotrophic industrial hosts into autotrophic strains capable of assimilating CO₂ as their sole carbon source. For example, *Komagataella phaffii* and *Escherichia coli* have been engineered to support fully autotrophic growth through the introduction of the Calvin–Benson–Bassham (CBB) cycle, with cellular energy supplied by methanol and formate oxidation, respectively^10,11^. Adaptive lab evolution (ALE) was used to improve the growth rates of each organism, and in both cases enabling mutations related to energy metabolism were uncovered^12,13^. Autotrophic *K. phaffii* strains were further developed to produce the industrial biopolymer monomers itaconic acid and lactic acid from CO_2_ with titers of 2 g L^-1^ and 0.6 g L^-1^L respectively^14^. Recently, by further strain and process optimisation, a titer of 12 g L^-1^ itaconic acid from CO_2_ could be achieved^15^. *K. phaffii* remains an interesting host for further development in the single carbon bioeconomy due to its ease of handling, genetic engineering tools, and robustness in industrial settings ^16–19^.

In addition to improving titers and production rates, a focus on improving the conversion efficiency of strains is important from both an economic and sustainability standpoint^20^. This consideration is particularly relevant in the context of the C1 bioeconomy, where high-volume products such as bulk chemicals and biomass are often the primary targets^21^. One promising strategy for improving process efficiency is the identification and activation of latent C1 metabolic pathways in industrial microorganisms^22–26^. In methylotrophic yeasts, the inefficient oxidation of methanol by alcohol oxidases (Aox) represents a key opportunity for intervention. Replacing this reaction with an alcohol dehydrogenase would generate an additional molecule of NADH per methanol consumed, thereby improving electron conservation and increasing overall substrate utilization efficiency^5^. It was recently discovered that *K. phaffii* strains lacking both *AOX1* and *AOX2* (so-called Mut⁻ strains) retain a basal capacity for methanol oxidation despite being unable to grow on methanol^27^. Through targeted gene knockouts and ^13^C-labeling experiments, the enzyme responsible for this residual methanol oxidation was identified as the native alcohol dehydrogenase 2 (Adh2)^28^. However, these studies did not achieve a growth phenotype in Mut⁻ strains with methanol as the sole carbon and energy source.

In this study, we explored the potential of the native Adh2 to support methanol dissimilation and improve carbon efficiency in synthetic autotrophic strains of *K. phaffii*. By deleting the native *AOX* genes, followed by *ADH2* overexpression, we demonstrate that Adh-based autotrophic strains can grow on CO₂ and methanol with improved carbon efficiency. In addition, we show production of both itaconic acid and lactic acid in Adh-based autotrophic strains, with marked improvements in carbon efficiency and productivity. Together, these results provide the first demonstration of *K. phaffii* growth on single-carbon substrates in the absence of alcohol oxidases and highlight the potential of Adh2-mediated methanol dissimilation for improving biomass formation and product yields.

## Results

### Strain engineering and theoretical improvements for Adh2-mediated methanol dissimilation

The autotrophic *K. phaffii* strain constructed by Gassler et al. (CO2-AOX) assimilates carbon into biomass from CO_2_ and relies on the dissimilation of methanol by alcohol oxidase 2 (Aox2) for energy generation ^10^. To engineer a strain dependent on Adh2-mediated methanol oxidation, *AOX2* was first knocked out to produce a Mut^-^ autotrophic strain of *K. phaffii* (CO2-Mut-). For additional flux through the MeOH oxidation reaction, a second copy of *ADH2* was overexpressed to produce the strain CO2-ADH (Fig. 1A). The Adh2 reaction reduces NAD^+^ to NADH during the oxidation on methanol, increasing the energetic yield of the dissimilatory pathway by 50% from 2 NADH to 3 NADH. For the conversion of 3 CO_2_ into 1 pyruvate via the synthetic CBB cycle, the theoretical energy requirement is 7 ATP and 5 NADH, equivalent to 19.5 ATP when assuming 2.5 ATP per NADH through the malate-aspartate shuttle. Based on the required methanol to meet the ATP demand, mass balance calculations predict a 33% reduction in methanol consumption per carbon fixed into pyruvate, decreasing from 1.30 to 0.87 mol mol⁻¹. The net CO₂ balance per mol carbon fixed into pyruvate shifts from a net production of +0.9 mol in CO2-AOX to a net consumption of −0.4 mol in CO2-ADH, enabling theoretical net fixation. However, this analysis does not account for additional CO₂ generation associated with cellular maintenance energy requirements.

**Fig. 1.**
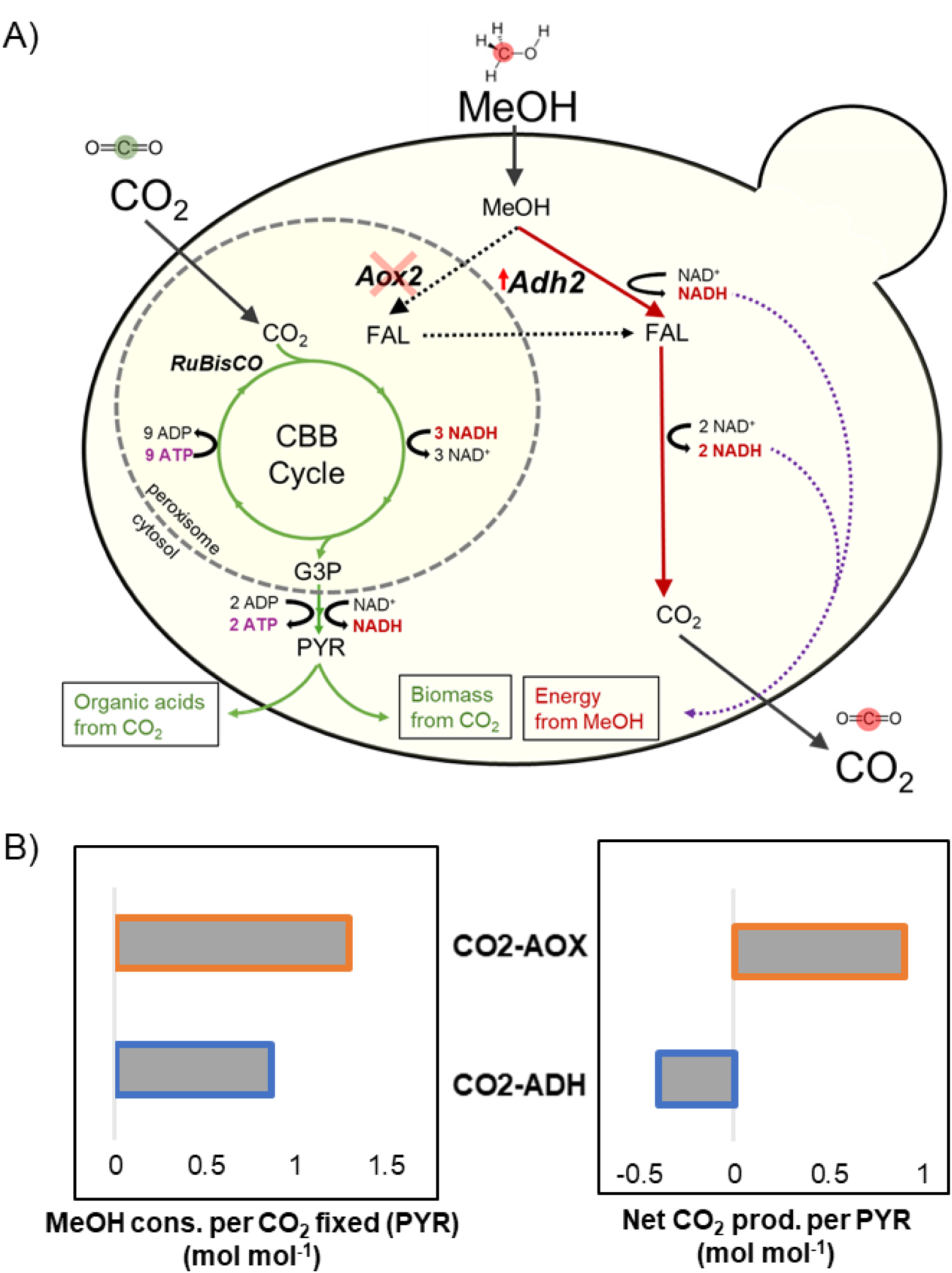
Schematic overview of carbon and energy metabolism in autotrophic *K. phaffii* with Adh-mediated dissimilation. A) CO2 is assimilated via an engineered Calvin-Benson-Bassham (CBB) cycle, requiring 9 ATP and 3 NADH per glyceraldehyde 3-phosphate (G3P). There is a recovery of 2 ATP and 1 NADH in the conversion of G3P to pyruvate (PYR), which can be incorporated into biomass or other (secreted) organic acids (in production strains). Energy is supplied through the dissimilatory pathway, where methanol is oxidized to CO2 producing either 2 NADH in CO2-AOX strains, or 3 NADH in the engineered CO2-ADH strain depicted here B) Comparison of theoretical efficiency of conversion of CO2 to pyruvate in synthetic autotrophic strains. A conversion of 2.5 ATP per NADH is used assuming use of the malate-aspartate shuttle for electron transfer into mitochondria. Full stoichiometric calculations are given in Table S1. *AOX2*, alcohol oxidase 2; *ADH2*, alcohol dehydrogenase 2; MeOH, methanol; FAL, formaldehyde; G3P, glyceraldehyde 3-phosphate; PYR, pyruvate.

### Overexpression of *ADH2* supports growth of Mut^-^ autotrophic strain

To compare growth of CO2-AOX and CO2-ADH, a shake flask experiment was performed in a CO_2_ shaking incubator at 30°C where 5% CO_2_ was supplied in the headspace, and the methanol concentration in the medium was maintained at 1% vol vol^-1^ following each sampling. Over two weeks of cultivation, the strain CO2-Mut- exhibited minimal growth with an average growth rate (μ_avg_) between 4 h and 310 h of 0.0028 ± 0.0001 h^-1^. In contrast the strain with additional Adh2 overexpression, CO2-ADH, grew with a μ_avg._ of 0.0054 ± 0.0003 h^-1^, compared to 0.0092 ± 0.0002 h^-1^ for the Aox-based strain CO2-AOX (Fig. 2). To assess whether peroxisomal targeting of Adh2 would be beneficial, the CO2-ADH-Pts strain was constructed; however, this also exhibited only minimal growth (μ_avg =_ 0.0032 ± 0.0006 h^-1^). Together, these results indicate that cytosolic overexpression of *ADH2* is both necessary and sufficient to supply energy for autotrophic growth of Mut^-^ synthetic autotrophic *K. phaffii*.

**Fig. 2.**
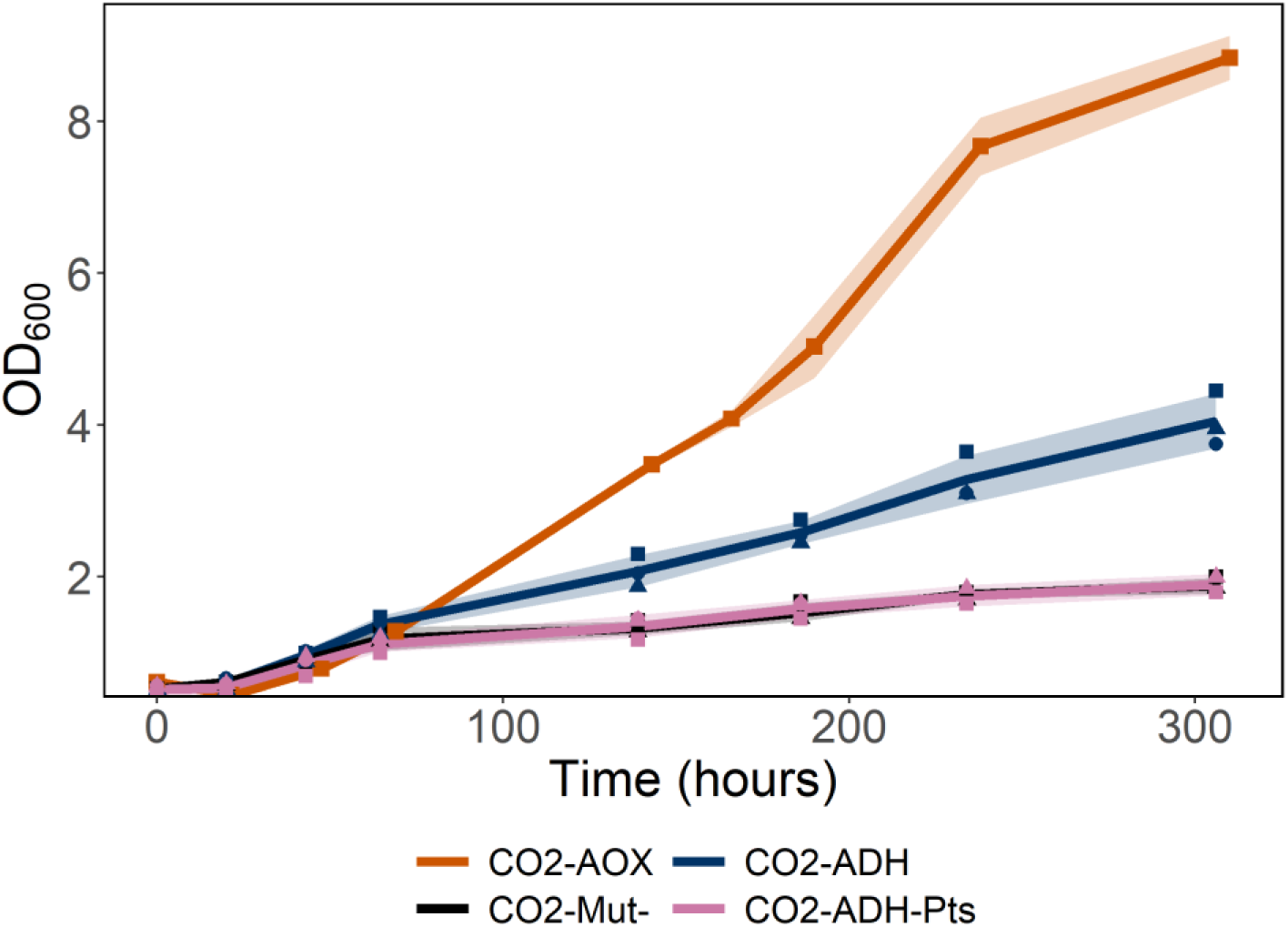
Cytosolic overexpression of *ADH2* supports growth in Mut- autotrophic *K. phaffii*. Cultivation of autotrophic strains engineered for Adh-mediated methanol oxidation. Strains were grown in shake flasks in a CO2 shaking-incubator at 30°C with 5% CO2 and 1% methanol supplied. For CO2-AOX (orange line), the mean of three technical replicates (individual cultivations) is shown; for CO2-Mut- (black line), CO2-ADH-pts (pink line) and CO2-ADH (blue line), the mean of three biological replicates (different clones in individual cultivations) is shown. The cultivations for CO2-AOX were performed in a separate experiment under identical conditions.

### Adh-based strains emit less CO_2_ and consume less methanol

With growth established, we next compared the carbon efficiency of the CO2-AOX and CO2-ADH strains. Cultivations were performed in a Transfer Rate Online Measurement (TOM) shaker system, in which each shake flask is equipped with a dedicated cap enabling individual, real-time monitoring of CO₂ transfer rate (CTR) and oxygen transfer rate (OTR) throughout the experiment^29^. As all strains exhibited net positive evolution of CO_2_, we defined improved carbon efficiency as reduced CO₂ evolution relative to biomass formation. At 30 °C, both strains displayed similar growth in the TOM-shaker unit (average μ = 0.0065 vs 0.0070), while CO2-ADH exhibited lower specific CO₂ evolution (q_CO₂_) and specific oxygen consumption (q_O₂_) over the course of cultivation (Table 1., Fig. 3A–C). Averaged across the experiment, q_O2_ and q_CO2_ were reduced by 46% and 50%, respectively, for CO2-ADH compared with CO2-AOX (Table 1). To quantify carbon efficiency on a molar basis, cumulative carbon incorporation into biomass and release as CO₂ were calculated over the duration of the TOM-shaker experiment (Table S2). The total moles of CO₂ evolved were evaluated relative to the total moles of carbon fixed into biomass to determine the molar carbon yield, Y_X/CO₂_ (C-mol C- mol⁻¹), which was improved 2.1-fold in CO2-ADH compared with CO2-AOX (Fig. 3C).

**Fig. 3.**
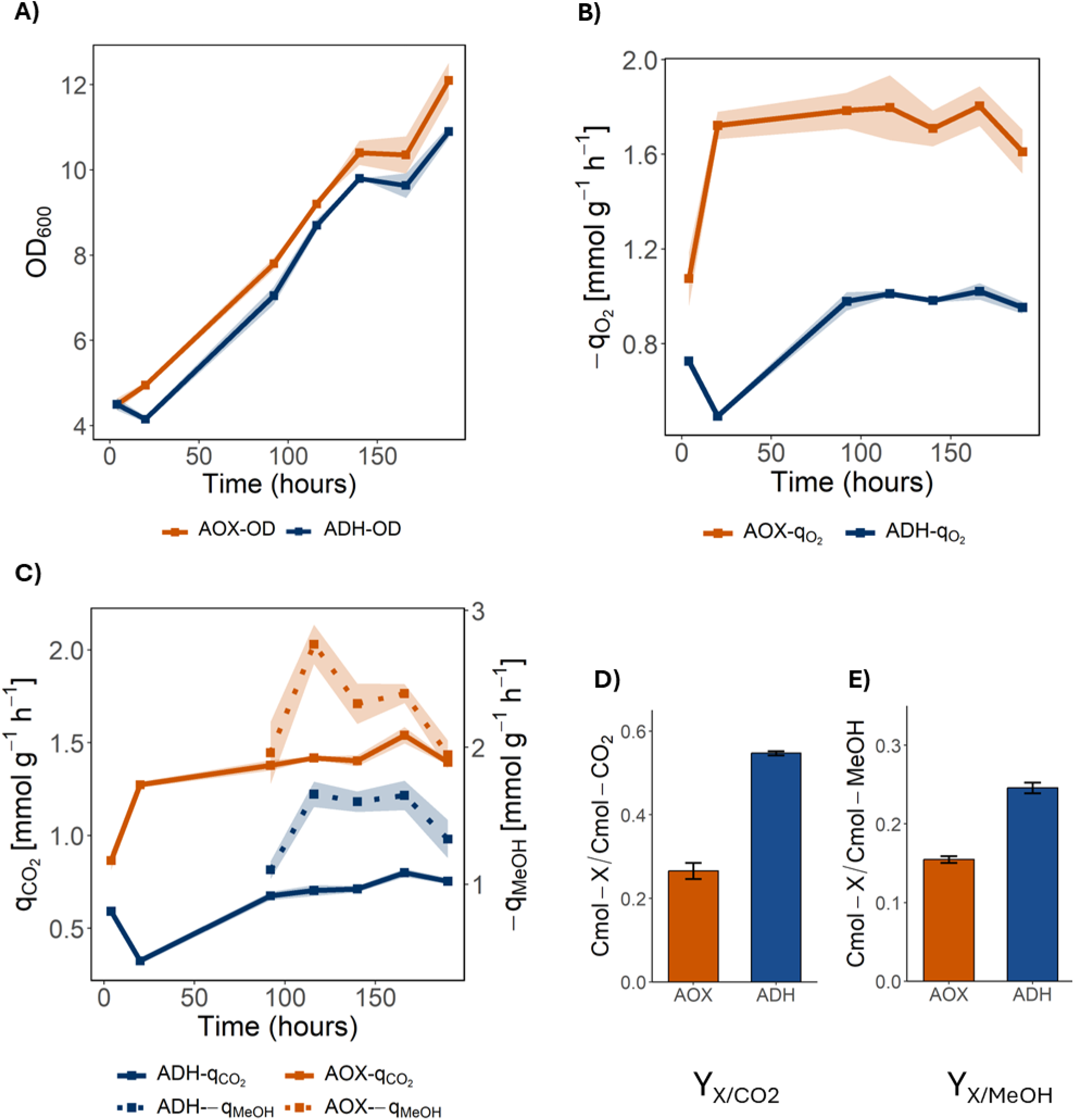
Comparison of growth, methanol consumption, and net CO2 evolution for Adh-based and Aox-based non-producing strains. Cultivations performed in TOM-shaker with analysis of (A) optical density, (B) specific oxygen uptake (qO2), (C) specific CO2 evolution (qCO2) and specific methanol consumption (qMeOH). Total mol carbon fixed into biomass was assessed in relation to (D) total net mol CO2 evolved (YX/CO2) and (E) total mol methanol consumed (YX/MeOH) The orange lines and columns represents CO2-AOX and blue represents CO2-ADH. The mean of two individual cultivations for each strain is shown. Standard deviation (±) is represented by either shading (line plots) or error bars (column plots).

**Table 1.** Average values of key process efficiency parameters.

| Strain | MeOH target<br>[vol vol<sup>-1</sup>] | temp.<br>[°C] | $\mu$<br>[h<sup>-1</sup>] | Average values | | | |
| --- | --- | --- | --- | --- | --- | --- | --- |
| | | | | $Y_{x/s}$<br>[g g<sup>-1</sup>] | $q_{MeOH}$<br>[mmol·g<sup>-1</sup>·h<sup>-1</sup>] | $q_{CO_2}$<br>[mmol·g<sup>-1</sup>·h<sup>-1</sup>] | $q_{O_2}$<br>[mmol·g<sup>-1</sup>·h<sup>-1</sup>] |
| CO<sub>2</sub>-AOX | 1% | 30 | 0.0070 | 0.16 | -1.27 | 0.81 | -1.71 |
| CO<sub>2</sub>-ADH | 1% | 30 | 0.0065 | 0.24 | -0.83 | 0.40 | -0.92 |

To quantify methanol consumption of CO2-AOX and CO2-ADH strains, methanol loss due to evaporation during TOM-shaker cultivation was first determined using cell-free control flasks containing medium only, sampled in parallel with experimental cultures. Throughout the experiment the specific methanol consumption (q_MeOH_) remained lower for CO2-ADH strains while maintaining a similar growth rate (Fig. 3A, C, Table 1). Cumulative methanol utilization was subsequently used to calculate biomass yield over the full cultivation period. The biomass yield Y_X/S_ (g g^-1^) increased by 59%, and qMeOH was reduced by 35%, for CO2-ADH relative to CO2-AOX (Table 1). The molar biomass yield, Y_X/MeOH_ (Cmol Cmol^-1^), was also determined, revealing a 60% improvement for CO2-ADH compared with CO2-AOX (Fig. 3D).

### Reduced oxygen supply leads to similar growth for Aox- and Adh-based strains

After observing improved performance of Adh-based autotrophic strains in TOM-shaker cultivations compared with conventional shake flasks (Fig. 2, Fig. 3A), we hypothesized that this effect may be related to reduced oxygen availability resulting from the TOM-shaker caps used for gas analysis, as well as cycling between aeration and measurement phases ^29^. To further investigate the impact of oxygen limitation on Adh- and Aox-based strains, bioreactor cultivations were performed with the inlet oxygen concentration reduced to 10%. Under these conditions, both strains exhibited similar overall growth profiles, consistent with the TOM-shaker experiments (Fig. 4A). The large divergence in dissolved oxygen and CO_2_ evolution rate (CER) profiles further indicated reduced oxygen consumption and improved carbon efficiency of CO2-ADH. To quantify carbon efficiency improvements in this context, the cumulative molar carbon yield Y_X/CO₂_ (mol mol⁻¹) was calculated (Fig. 4D), revealing a 1.86-fold improvement for CO₂-ADH compared with CO₂-AOX. This result closely aligns with the 2.1-fold increase observed for the same strains in the TOM-shaker experiments.

**Fig. 4.**
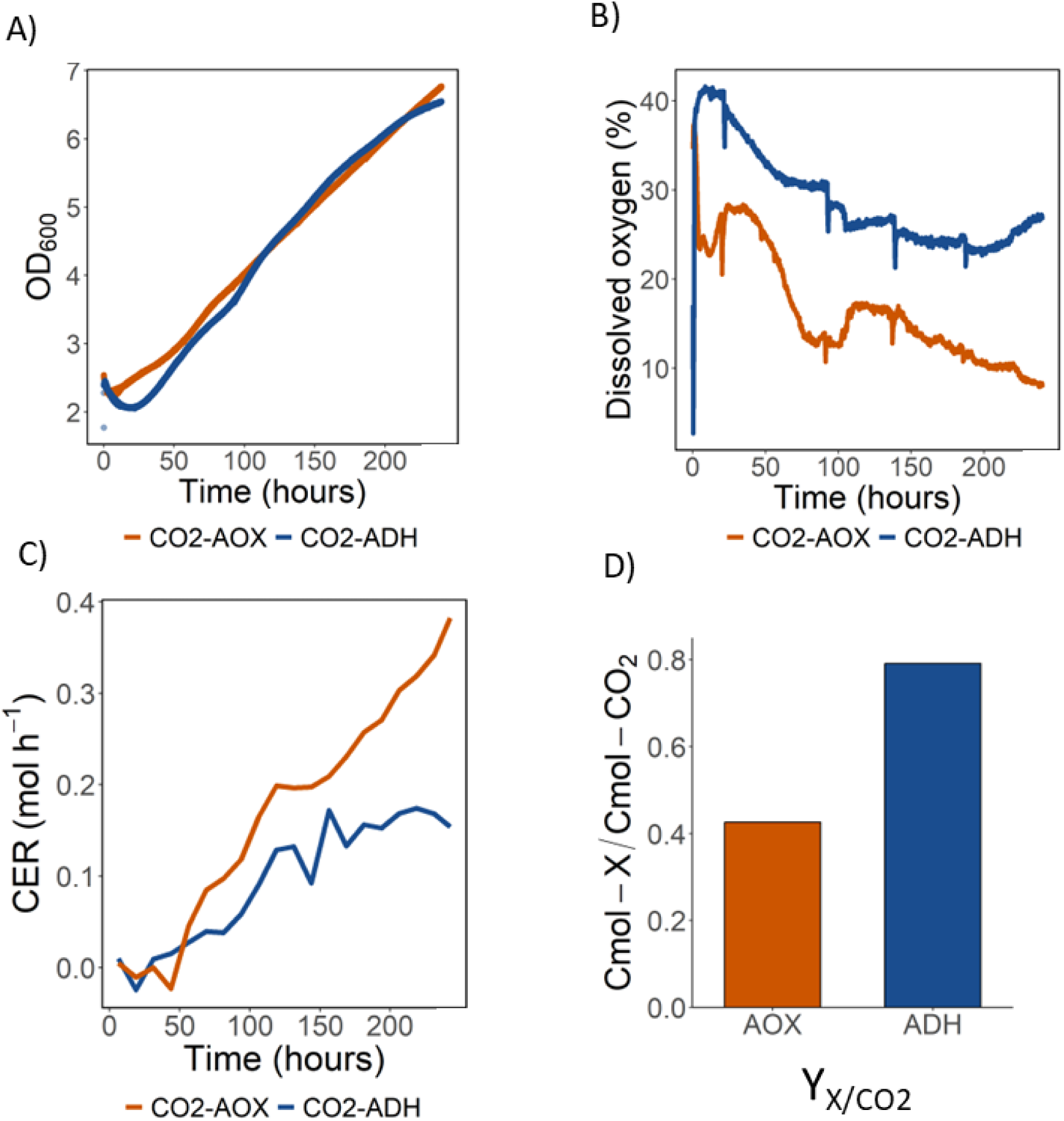
Bioreactor cultivation of CO2-AOX and CO2-ADH strains in oxygen-limited conditions. Online measurements of (A) growth, (B) dissolved oxygen, and (C) CO2 evolution rate (CER) are shown. D) molar biomass yield (YX/CO2) for non-producing CO2-AOX and CO2-ADH strains. Cultivations were performed at 30°C with 5% CO₂ and 10% oxygen supplied at the inlet.

### Adh2 supports superior organic acid production from CO_2_ in autotrophic strains

Baumschabl et. al recently engineered autotrophic strains of *K. phaffii* capable of converting CO_2_ into both itaconic and lactic acid via the synthetic CBB cycle, relying on Aox2 for methanol oxidation^14^. The itaconic acid production strain was used directly in this work (CO2-IA-AOX). The lactic acid production strain, which was engineered to reduce lactate re-consumption through deletion of the L-lactate cytochrome-c oxidoreductase (*CYB2*), was further modified for this work. Baumschabl et al. reported that RuBisCO oxygenation in autotrophic *K. phaffii* leads to glycolate formation, which is subsequently recycled via a native phosphoglycolate salvage pathway^22^. In *cyb2Δ* strains, however, glycolate accumulated to high levels, indicating that Cyb2 catalyzes oxidation of glycolate to glyoxylate. To prevent glycolate accumulation, glycolate dehydrogenase (*GYD1*) was overexpressed to promote oxidation of glycolate, generating the lactic acid base strain used in this study (CO2-LA-AOX).

Both CO2-LA-AOX and CO2-IA-AOX were converted to Adh2-dependent methanol utilization first by performing a CRISPR-Cas9-mediated replacement of the *AOX2* coding sequence with *ADH2,* yielding CO2-IA-Mut- + CO2-LA-Mut- strains. Multi-copy integration of P_AOX1__*ADH2* was then performed using a zeocin-resistance plasmid to generate strains with elevated *ADH2* gene copy number (GCN). A range of 1 - 3 additional copies of *ADH2* were attained for lactic acid strains, and 1 - 6 additional copies for itaconic acid production strains, respectively. A shake-flask experiment feeding CO_2_ and methanol was performed to assess multi-copy strains for production of organic acids from CO_2_. Lactic acid and itaconic acid concentrations were monitored over 10 days, and final titers were analysed as a function of *ADH2* gene copy number (GCN) (Fig. S1A, B). For lactic acid production, titers increased with increasing *ADH2* copy number, whereas for itaconic acid production, maximal titers were observed at an *ADH2* GCN of four, with higher copy numbers resulting in progressively lower titers. The best-performing strains from this screening, LA-MC5 and IA-MC2 (Fig. S1), were selected for further characterization and designated CO2-LA-ADH and CO2-IA-ADH, respectively.

Ata et al. reported that Aox-based autotrophic *K. phaffii* production strains performed optimally at 25°C, exhibiting higher titers and growth rates compared with cultivation at 30°C. In contrast, Adh-based autotrophic strains in this study showed improved performance at 30°C relative to 25°C, consistent with more favorable thermodynamics of the Adh reaction at elevated temperatures^30^. As methanol dehydrogenases are known to have poor affinity for methanol compared with alcohol oxidases, the effect of increased methanol concentrations on CO2-ADH strains was also tested to improve flux through the Adh2 reaction^31^. Increasing methanol concentration from 1% to 2% (vol vol^-1^) improved lactic acid production for CO2-LA-ADH, resulting in an increase of the final titer from 650 mg L^-1^ to 950 mg L^-1^ while slightly reducing biomass formation (Fig. S1C). A further increase from 2% to 3% (vol vol^-1^) led to a modest additional improvement in lactate titer, from 950 mg L⁻¹ to 1100 mg L⁻¹. For itaconate production, elevating the methanol concentration target from the beginning of the cultivation resulted in reduced final titers for Adh-based strains, with little effect on biomass formation (Fig. S1D).

### Adh-based strains maintain higher organic acid productivity with reduced CO_2_ evolution

With growth and organic acid production established, we next compared the carbon efficiency of Adh-based and Aox-based production strains. Extending stoichiometric calculations from Fig. 1 to include the energy demands for conversion of pyruvate into either itaconic acid or lactic acid predicts methanol requirements of 1.50 and 1.47 mol MeOH per mol CO₂ fixed into product for CO2-IA-AOX and CO2-LA-AOX, respectively (Fig. 5, Table S3). In contrast, the corresponding methanol requirements for CO2-IA-ADH and CO2-LA-ADH were reduced to 1.00 and 0.98 mol MeOH per mol CO₂ fixed into product, respectively. Due to the decarboxylation reaction catalysed by CadA, theoretical net CO_2_ production for itaconic acid is higher than for lactic acid. In both cases, net CO₂ evolution was reduced in Adh-based strains, with zero net CO₂ production predicted for CO2-IA-ADH and slight net fixation predicted for CO2-LA-ADH. However, these calculations do not account for cellular maintenance energy requirements.

**Fig. 5.**
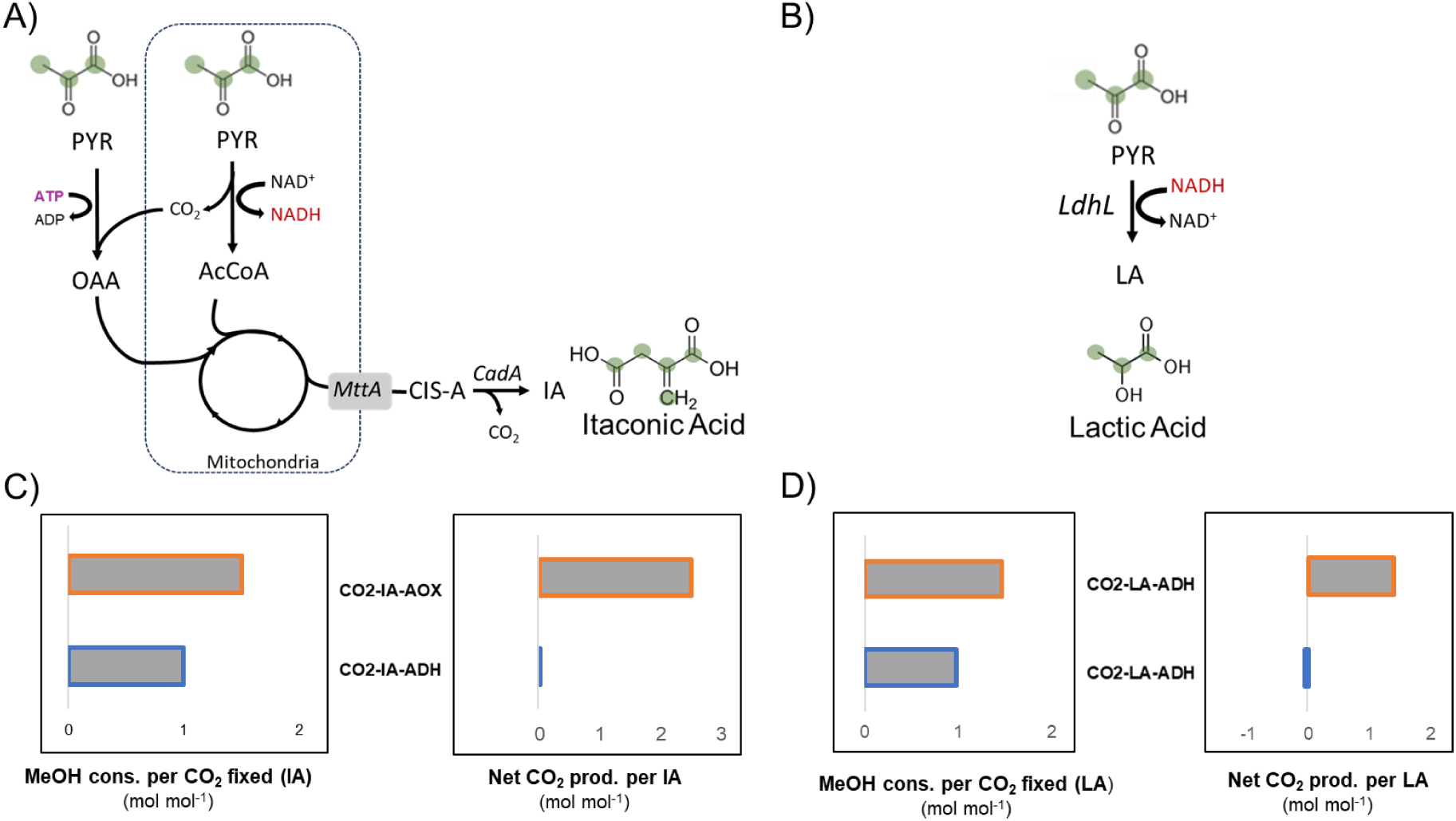
Schematic overview of carbon and energy metabolism for production of organic acids from of pyruvate in engineered autotrophic *K. phaffii* strains. A) Conversion of pyruvate to itaconic acid in CO2-IA-AOX and CO2-LA-AOX strains: 1 itaconic acid (IA) is produced from 2 pyruvate (PYR) with concomitant release of 2 CO2 + 1 NADH and consumption of 1 ATP + 1 CO2. B) Conversion of pyruvate to lactic acid in CO2-LA-AOX and CO2-LA-ADH strains: 1 lactic acid is produced from 1 pyruvate (LA) with consumption of 1 NADH (C) Comparison of theoretical efficiency of CO2-to-itaconic acid conversion in CO2-IA-AOX versus CO2-IA-ADH. (D) Comparison of theoretical efficiency of CO2-to-lactic acid conversion in CO2-LA-AOX versus CO2-LA-ADH. An ATP yield of 2.5 per NADH was assumed. Full stoichiometric calculations are given in Table S3. Cons. = consumed; prod. = produced. PYR, pyruvate; AcCoA, acetyl-coA; CIS-A, cis-iconitate; OAA, oxaloacetate; IA, itaconic acid; LA, lactic acid. Enzymatic steps catalysed by heterologous proteins are indicated in italics: *MttA*, mitochondrial tricarboxylic acid transporter; *CadA*, cis-aconitate decarboxylase; *LdhL*, lactate dehydrogenase.

To assess experimental carbon efficiency of production strains, cultivations were performed in TOM-shaker incubator units with online monitoring of CTR and OTR. As all production strains were net producers of CO_2_, improved carbon efficiency was again defined as reduced net CO_2_ evolution relative to biomass and organic acid formation. At 30°C and 1% methanol (vol vol^-1^) both CO2-ADH strains exhibited higher specific productivities (q_LA,_ q_IA_) and growth rates while maintaining lower q_CO2_ compared to CO2-AOX strains (Fig. 6A,B). Relative to CO2-LA-AOX, the average growth rate (μ_avg_) and q_P_ were increased for CO_2_-LA-ADH by 25% and 4.4-fold, respectively, while the net q_CO2_ was reduced by 16% (Table 2). Similarly, compared with CO_2_-IA-AOX strain, CO_2_-IA-ADH had more than 80-fold increase in growth rate and a 35% increase in q_P_, together with a 20% reduction in q_CO2_.

**Fig. 6.**
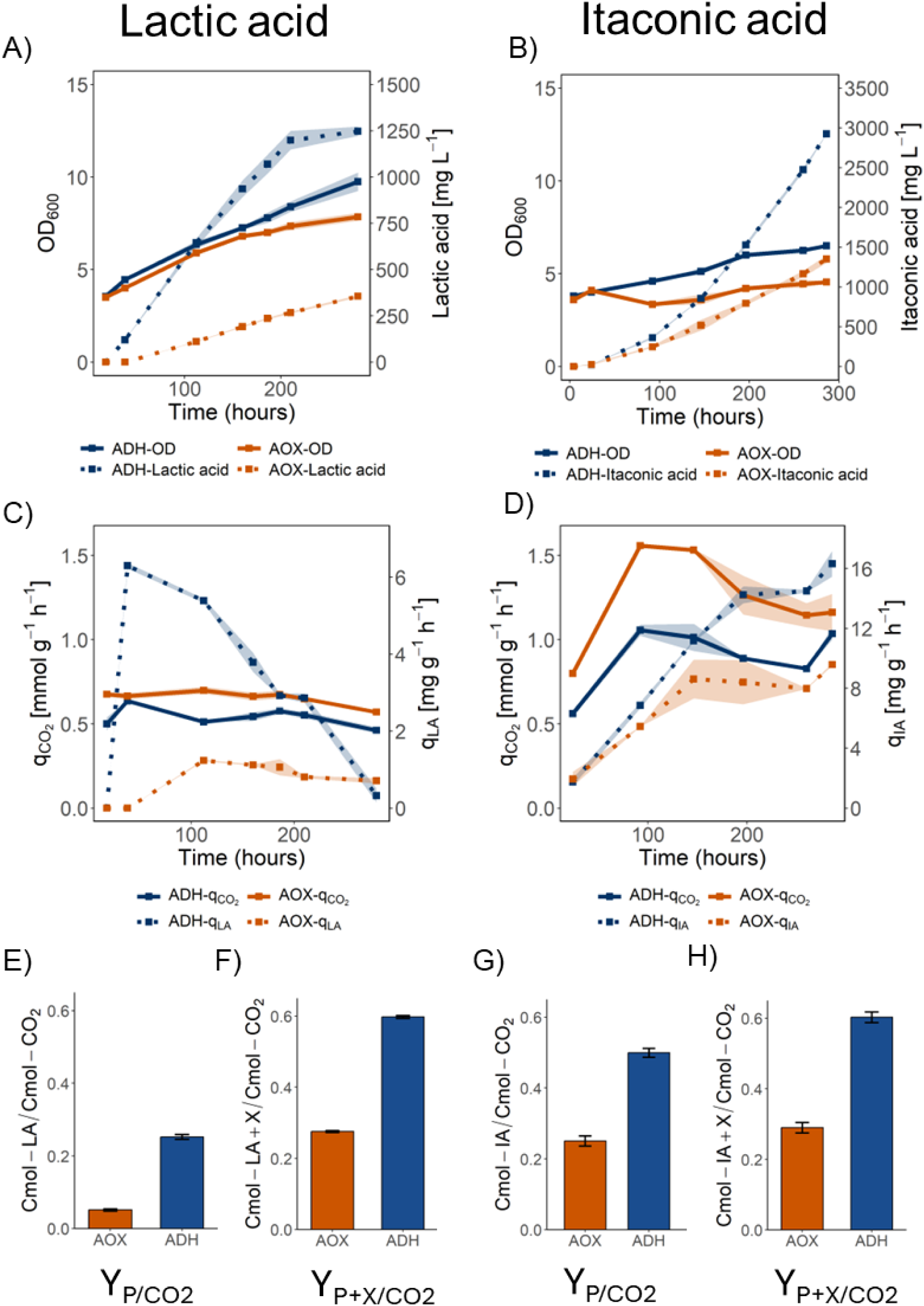
Comparison of growth, organic acid production, and net CO2 evolution for Adh- and Aox-based production strains. Cultivations of CO2-LA-AOX, CO2-LA-ADH, CO2-IA-AOX and CO2-IA-ADH strains performed in TOM-shaker units at 30°C with 5% CO2 and 1% methanol (vol vol^-1^) supplied. (A, B) Optical density and product titer, and (C,D) specific CO2 evolution rate (qCO2) and specific productivity (qLA, qIA). Total mol carbon evolved as CO2 assessed in relation to (E) total mol carbon fixed as lactic acid (P), (F) total mol carbon fixed as lactic acid biomass (X), (G) total mol carbon fixed as itaconic acid (P), and (H) total mol carbon fixed as itaconic acid and biomass (X).Orange lines and columns indicate Aox-based strains (CO2-LA-AOX, CO2- IA-AOX), while blue lines and bars indicate Adh-based strains (CO2-LA-ADH, CO2-IA-ADH). The mean of two individual cultivations for each strain is shown. Standard deviation (±) is represented by either shading (line plots) or error bars (column plots).

**Table 2.** Average values of key parameters for production strains.

| Strain | MeOH target<br>[vol vol<sup>-1</sup>] | Temp<br>[°C] | $\mu$<br>[h<sup>-1</sup>] | $q_p$<br>[mg·g<sup>-1</sup>·h<sup>-1</sup>] | $q_{CO_2}$<br>[mmol·g<sup>-1</sup>·h<sup>-1</sup>] | $q_{O_2}$<br>[mmol·g<sup>-1</sup>·h<sup>-1</sup>] |
| --- | --- | --- | --- | --- | --- | --- |
| CO2-IA-AOX | 1% | 30 | 0.0004 | 5.25 | 0.955 | -2.19 |
| CO2-IA-ADH | 1% | 30 | 0.0033 | 8.12 | 0.675 | -1.66 |
| CO2-IA-ADH | 1.5% | 30 | 0.0037 | 8.67 | 0.751 | -1.76 |
| CO2-LA-AOX | 1% | 30 | 0.0053 | 0.82 | 0.654 | -1.12 |
| CO2-LA-ADH | 1% | 30 | 0.0067 | 3.60 | 0.547 | -1.21 |
| CO2-LA-ADH | 2% | 30 | 0.0059 | 4.30 | 0.648 | -1.38 |

For CO2-LA-ADH, increasing the target methanol concentration resulted in higher lactate production accompanied by increased CO₂ evolution (Table 2). With a 2% methanol (vol vol^-1^) target, specific productivity (q_P_) and q_CO2_ were increased 19% and 18% respectively, compared with 1% (vol vol^-1^). In addition, at 2% methanol (vol vol^-1^) the strain continued producing throughout the two-week cultivation, rather than reaching a plateau after approximately 200 h (Fig. S2). Elevated methanol concentration was also tested for CO2-LA-AOX strains; however, this led to further reductions in growth and production rates (Table 2, Fig. S2). A modest improvement in productivity was observed for CO2-IA-ADH strains when the methanol concentration target was increased mid-way through the experiment (Table 2, Fig. S2). Compared with a 1% methanol (vol vol^-1^) target throughout the whole experiment, raising to 1.5% vol vol^-1^ on day 5 resulted in an increase in q_P_ and q_CO2_ for CO2-IA-ADH by 6 % and 11 %, respectively. Consistent with data for non-production strains, reduced q_O2_ was also observed for CO2-IA-ADH, which exhibited a 24% lower q_O2_ than CO2-IA-AOX in the same conditions despite producing higher levels of both biomass and product (Table 2). In contrast, during cultivation at 1% methanol (vol vol^-1^), the average q_O2_ for CO2-LA-ADH was 8% higher compared with CO2-LA-AOX, consistent with a larger increase in q_P_ for Adh-based lactic acid-producing strains.

To quantify conversion efficiency on a molar basis, total carbon incorporated into biomass and product, as well as carbon released as CO₂, was determined for the TOM-shaker cultivations (Table S6). Cumulative carbon evolved as CO_2_ over the course of each experiment was examined in relation to the total carbon fixed into product and biomass (Fig. 6 E-H). First, the mol carbon evolved was compared to the carbon fixed solely into product (Y_P/CO2_). Relative to Aox-based strains, CO2-LA-ADH and CO2-IA-ADH exhibited 4-fold and 2-fold increases in Y_P/CO_₂, respectively (Fig. 6E, G). The carbon produced as biomass (Cmol X) was then also included to assess total carbon fixation relative to net CO_2_ evolution (Y_(X+P)/CO2_) (Fig. 6 F,H). The 2-fold improvement in Y_(X+P)/CO2_ (Cmol Cmol^-1^) of Adh-based strains for lactic acid production then matched closely with that of itaconic acid production strains.

Aox-based strains were also evaluated for carbon efficiency in the TOM-shaker at 25°C, a temperature previously shown to support higher growth rates and product formation^15^. In these conditions, all strains had higher growth rates and q_CO2_ than at 30°C (Fig. S2, Table S5). For CO2-IA-AOX, the itaconic acid titer reached 2876 mg L⁻¹ at 25°C, representing more than a 2-fold increase relative to 30°C; however, this value remained lower than that achieved by CO2-IA-ADH at 30°C. In addition, owing to substantially increased biomass formation, specific productivity (q_P_) was markedly reduced for CO2-IA-AOX at 25°C (Table S5). For CO2-LA-AOX, the initial productivity was considerably higher at 25°C, exceeding 350 mg L^-1^ at hour 140, after which re-consumption was observed. Molar carbon yields were improved for all Aox strains at 25°C by roughly 40% compared to 30°C, yet remained roughly 50% lower than for Adh strains at 30°C. Together, these results indicate that while reduced temperature improves growth and product titers in Aox-based strains, both their carbon efficiency and productivity remain lower than that of Adh-based strains cultivated at 30°C.

### Lactic acid production from CO_2_ in Adh-based strains is improved by further strain engineering

It was recently reported that lactic acid represses methanol utilization genes in *K. phaffii*, and that overexpression of the native genes *MIT1* and *MXR1,* coding for transcription factors involved in methanol metabolism, can help maintain higher lactate production from methanol^32^. A Multigene construct containing overexpression cassettes of both *MIT1* and *MXR1* was inserted into the strain CO2-LA-ADH (CO2-LA-ADH_Mxr-Mit), leading to an increase in titer of 25% at 140 h (Fig. S4). Due to a delay of the reduction in lactic acid productivity, a maximum titer of 1600 mg L^-1^ lactic acid was also reached, representing an 50% increase in titer relative to the parental CO2-LA-ADH strain in the same conditions. Cultivation of CO2-LA-ADH_Mxr-Mit at higher initial cell density (OD = 8) further increased production, yielding more than 1700 mg L⁻¹ lactic acid within 140 h, although product consumption was still observed at later time points.

## Discussion

In this study, we explored the potential of Adh2-mediated methanol oxidation to improve the carbon efficiency of growth and chemical production in autotrophic *K. phaffii* strains. To date, all methylotrophic yeasts described in the literature, including members of the genera *Komagataella*, *Ogataea*, and *Candida*, have been reported to rely on alcohol oxidases for methanol oxidation^33^. Therefore, the growth phenotype observed for the CO2-ADH strain represents, to our knowledge, the first example of a methylotrophic yeast capable of growth on single-carbon substrates in the absence of alcohol oxidases. Zavec et al. identified Adh2-mediated methanol utilization in Mut^-^ *K. phaffii* strains; however, in these strains close to 100% of the methanol was dissimilated to CO_2_, and growth was not achieved^28^. The decoupling of assimilation and dissimilation in autotrophic *K. phaffii* strains is likely the factor allowing for Adh-mediated growth in the present study (Fig. 1A). Because carbon assimilation is supplied from CO₂ via the CBB cycle, formaldehyde is not required for assimilatory metabolism, thereby eliminating competition between assimilatory and dissimilatory pathways, which may have limited growth in methylotrophic Mut⁻ strains. Moreover, unlike in methylotrophic Mut^-^ (Adh-based) strains, the cytosolic localization of formaldehyde production in this context is advantageous, as it is co-localized with the rest of the dissimilatory pathway. Nevertheless, the reduced growth rate of CO2-ADH relative to CO2-AOX observed in initial shake-flask experiments was expected, given the comparatively low activity of alcohol dehydrogenases toward methanol (Fig. 2)^31^. The improved performance of Adh-based strains at higher temperatures is also in alignment with thermodynamic predictions and experimental data from methanol dehydrogenases^30^.

Comparison of the non-producing CO2-AOX and CO2-ADH strains in TOM-shaker cultivations provides clear evidence for improved carbon efficiency of autotrophic growth with Adh-mediated methanol dissimilation (Fig. 3). In the same conditions and at a similar growth rate, q_CO2_ decreased by approximately 50%, while Y_X/S_ was increased by 60% in CO2-ADH. These results indicate that NADH generated through the Adh2-catalysed reaction is effectively incorporated into cellular energy metabolism, thereby reducing the amount of methanol required to be dissimilated to CO_2_ (Fig. 1 A), which is supported by calculations based on stoichiometry (Fig. 1B). A predictive challenge for carbon efficiency arises from RuBisCO oxygenation, which generates 2-phosphoglycolate that must be recycled through an energy-intensive salvage pathway. Baumschabl et al. demonstrated that this oxygenation reaction occurs at appreciable levels in autotrophic *K. phaffii* and is influenced by factors such as dissolved oxygen^22^. The reduced oxygen consumption observed for the CO2-ADH strain is also in alignment with stoichiometric calculations, as replacement of alcohol oxidase with Adh eliminates the requirement for approximately 1.5 mol O₂ per pyruvate produced from CO₂. This effect was evident both in TOM-shaker experiments, where lower q_O₂_ was observed throughout cultivation, and in bioreactor experiments, where dissolved oxygen declined substantially faster in CO2-AOX than in CO2-ADH (Table 1, Fig. 4B).

Classens et al. compared experimental yield data from a range of autotrophic microbial species to benchmark conversion efficiencies with respect to their energy source^34^. The Y_X/S_ calculated here for CO2-AOX (0.16 g g^-1^) is only slightly lower than the reported range of natural autotrophic bacteria which grow via the CBB cycle and utilize methanol as energy source (0.19 – 0.26 g g^-1^) (compiled by ^34^). In contrast, the Y_X/S_ of 0.24 g g^-1^ calculated for CO2-ADH approaches the upper end of this range, although it remains below than reported values for *methylotrophic yeasts* growing via the XuMP cycle (0.29-0.43 g g^-1^)(compiled by ^34^). This is consistent with flux balance analysis predictions for *E. coli* growing on single-carbon substrates through different metabolic routes, which indicate that CBB cycle–based growth results in lower biomass yields than growth via either the XuMP (syn. DHA) cycle or reductive glycine pathways^35^. The higher yield observed for the CO2-AOX strain growing at 25°C compared to 30°C could be due to the effect of increased growth rate (Table S5)^36^; however, this is difficult to separate from other temperature-dependent factors, including dissolved CO_2_ availability and RuBisCO selectivity.

Although net CO₂ evolution is often not reported for engineered autotrophic microbial strains, our findings suggest that most autotrophs relying on formate or methanol consumption for energy are likely to be net CO₂ producers. Therefore, this is an important process parameter to consider alongside biomass or product yield, as it impacts the carbon footprint of the overall process^20^. While q_CO2_ changes with growth and production rates, we find that the molar carbon yield Y_X/CO2_ provides an effective way to compare carbon efficiencies of different strains in batch cultivations. This can be particularly useful for methanol-based processes, where significant evaporation complicates accurate substrate-based yield calculations. While the absolute values Y_X/CO2_ for CO2-AOX and CO2-ADH differed between the bioreactor and TOM-shaker experiments, the relative improvement observed for CO2-ADH was highly consistent across both platforms. In addition to differences in cultivation equipment and analysis, part of this variation may be attributed to oxygen availability, with 10% oxygen supplied in the bioreactor compared with 21% in the TOM-shaker system. Additionally, 10% CO2 was supplied in the bioreactor experiment versus 5% for the TOM-shaker experiments.

We further demonstrated the improved carbon efficiency of Adh-based strains through comparison of autotrophic organic acid production strains. Relative to their Aox-based counterparts under identical conditions, both CO2-LA-ADH and CO2-IA-ADH strains exhibited lower q_CO2_ while maintaining higher specific growth rates and q_P_ (Fig. 6A, B). This provides clear evidence of enhanced metabolic efficiency in Adh-based strains, as increased assimilatory flux associated with higher biomass and product formation would be expected to require additional energy generation via methanol oxidation to CO₂. This relationship was observed for CO2-LA-ADH strains cultivated at 2% methanol (vol vol^-1^), where q_CO2_ increased roughly in proportion to q_P_ relative to cultivation at 1% methanol (vol vol^-1^) (Table 2). Carbon efficiency of Adh- and Aox-based production strains was also quantified using molar carbon yields, Y_P/CO2_ and Y_(P+X)/CO2_ (Fig. 6E-H). While substantial improvements in Y_P/CO2_ were observed for both product targets, lactic acid production efficiency particularly benefited from Adh2-mediated methanol dissimilation. Incorporating also biomass production, Y_(P+X)/CO2_ for both itaconic acid and lactic acid had approximately a 2-fold improvement for Adh2-based strains, converging to similar values of ∼0.6 C-mol C-mol⁻¹. For Aox-based production strains cultivated at 25°C, Y_P/CO2_ was lower due to high biomass formation, but Y_(P+X)/CO2_ improved for CO2-IA-AOX and CO2-LA-AOX by 32% and 39% respectively, compared with production at 30°C. However, in both cases this was still roughly 64% less than Adh-based counterparts at 30°C. Differences in gas solubility, RuBisCO selectivity, and growth rate may be contributing factors to the observed differences. Therefore, we consider comparisons performed under identical conditions (30°C, 1% MeOH) and similar growth rates to provide the most appropriate assessment of the relative carbon efficiencies of strains with Aox- versus Adh-mediated methanol oxidation.

Although improved efficiency was anticipated and was the primary focus of the study, the magnitude of the associated gains in productivity was unexpected. Particularly in the case of lactic acid production, the expanded cytosolic NADH pool in Adh-based strains appears to have promoted forward flux through the Ldh reaction, resulting in increased titers and a delayed onset of lactic acid re-consumption (Fig. 6A, S1–3). In addition to driving forward the Adh2 reaction, increased methanol concentrations may have conferred an benefit to CO2-LA-ADH strains by partially relieving lactate-mediated repression of methanol-regulated genes. However, CO2-LA-AOX did not benefit from higher methanol concentrations (Fig. S3), and was outperformed 4-fold by CO2-LA-ADH at the same methanol concentration (1% vol vol^-1^). Bachleitner et al. aimed to alleviate lactate-mediated repression through strain engineering and found that overexpression of the methanol-regulated transcription factors *MIT1* and *MXR1* in *K. phaffii* substantially increased final lactic acid titers from 4 to 17 g L⁻¹ and delayed the onset of lactic acid re-consumption in methanol-grown strains^32^. Similarly, in the present study an improvement in lactic acid productivity was also achieved with the *MIT1* + *MXR1* overexpression, reaching 50% higher titer than the parental CO2-LA-ADH strain (Fig. S4). Additionally, the strain CO2-LA-ADH_MIT-MXR generated very little biomass, which would improve the overall product yield and make it an even more attractive production host. To determine whether further increased titers could be achieved with this strain, a cultivation was performed at elevated cell density (starting OD=8). Under these conditions, a maximum lactic acid titer of 1.7 g L⁻¹ was reached after 140 h, representing a 60% increase compared to the same strain cultivated from a starting OD=4. To our knowledge, this represents the highest reported titer for lactic acid production from CO₂ in a microbial host. However, lactic acid re-consumption was still observed at later time points, resulting in a lower final titer than that obtained in the lower-cell-density cultivation. These results together indicate that increasing the cytosolic NADH pool is a promising strategy for improving lactic acid production in microbial production platforms; however, solutions to mitigate both lactic acid re-consumption and repression of methanol-regulated genes need further development.

The lack of improvement for Adh-based itaconate producing strains at higher *ADH2* gene copy numbers or elevated methanol concentrations may result from excess reducing equivalents entering the respiratory chain, which must then be dissipated^35^. This is supported by the greater increase in q_CO2_ production for CO2-IA-ADH at 1.5% MeOH (vol vol^-1^) relative to the increase in productivity. Nevertheless, with 4 copies of *ADH2*, CO2-IA-ADH achieved higher itaconic acid titers than the parental CO2-IA-AOX strains even at 25°C, while generating substantially less biomass (Table S4, Table S6). Ata et al. recently improved the itaconate titer of Aox-based autotrophic production strains to 12 g L^-1^ by optimizing copy numbers of genes involved in the CBB cycle and itaconate biosynthesis from the TCA cycle^15^. As this strategy targets the assimilatory branch, combining such approaches with Adh-mediated methanol dissimilation may provide complementary benefits, improving efficiency without compromising productivity. Moreover, that study highlights the importance on identifying the optimal process conditions for engineered autotrophic strains, which was not explored as extensively here. Further work to optimize initial cell density, CO_2_ supply, dissolved oxygen targets, and cultivation temperature is therefore likely to yield additional performance gains in Adh2-based autotrophic strains.

The key finding of this work is that the native Adh2 can substitute for Aox2 in methanol oxidation, enabling growth and chemical production in autotrophic *K. phaffii* strains with improved efficiency. A recent study similarly sought to increase the efficiency of *Cupriavidus necator* growing on CO_2_ and formate by replacing the native CBB cycle with a reductive glycine pathway, achieving a 17% increase in the Y_X/S_ (g g^-1^)^37^. In comparison, the 40% improvement in Y_X/S_ (g g^-1^) observed here underscores the strong potential of the Adh2- mediated methanol utilization as a strategy to improve efficiency of methylotrophic yeasts. However, even Adh-based autotrophic strains have a low biomass yield and high net CO_2_ production compared to methylotrophic *K. phaffii* strains, highlighting inefficiencies of C1 utilization based on the CBB cycle. In particular, reliance on RuBisCO for carbon fixation in the context of aerobic growth of yeasts presents challenges due to the oxygenation side reaction. These limitations may be partially mitigated through optimization of oxygen supply or implementation of CO₂-concentrating mechanisms such as carboxysomes, as employed by autotrophic bacteria^38,39^. Alternatively, pathways such as the CETCH cycle could be introduced, potentially enabling higher flux without the complication of an oxygenation side reaction; however, this synthetic cycle is less compatible with the native methanol metabolism of *K. phaffii* ^40^. Importantly, Adh-mediated methanol oxidation would be compatible as the energy generation module in the CETCH cycle, as well as other highly efficient C1 assimilation pathways already demonstrated in *K. phaffii*, including the XuMP cycle and the reductive glycine pathway^26^.

The primary objective of this study was to evaluate the potential for Adh-mediated methanol dissimilation to improve the carbon efficiency of autotrophic *K. phaffii* strains. Our results demonstrate that replacing Aox2 with Adh2 reduces CO_2_ production and increases biomass yields in *K. phaffii* strains growing via the CBB cycle. Elevated organic acid productivity was also observed for Adh-based strains, particularly for lactic acid production, where the additional reducing equivalents are directly utilized for product formation. Collectively, this work establishes Adh-mediated methanol oxidation as a broadly applicable strategy for engineering methylotrophic yeasts into more efficient microbial cell factories for converting single-carbon substrates into biomass and value-added chemicals.

## Materials and Methods

### Yeast strains

All *Komagataella phaffii* strains were based on the CBS7435 strain, and are summarised in Table 3.

**Table 3.** *K. phaffii* strains used in this study.

| Short name | Genotype | reference |
| --- | --- | --- |
| CO2-AOX | CBS7435 *aox1Δ::pAOX1_TDH3 + pFDH1_PRK + pALD4_PGK1 das1Δ::pDAS1_RuBisCO + pPDC1_groEL + pRPP1B_groES das2Δ::pDAS2_TLK1 + pRPS2_TPI1* | 10 |
| CO2-Mut- | CO2-AOX + *aox2Δ* | This study |
| CO2-ADH | CO2-AOX + *aox2Δ + pAOX1_ADH2* | This study |
| CO2-ADH-Pts | CO2-AOX + *aox2Δ + pAOX1_ADH2-PTS1* | This study |
| CO2-IA-AOX | CO2-AOX + *pAOX1_cadA + pPOR1_mttA* | 14 |
| CO2-IA-Mut- | CO2-IA-AOX + *aox2Δ::ADH2* | This study |
| CO2-IA-ADH | CO2-IA-Mut- + *pAOX1_ADH2* (multicopy clone – *ADH2* GCN=4) | This study |
| CO2-LA-AOX_parent | CO2-AOX + *pAOX1_ldhL + cyb2Δ* | 14 |
| CO2-LA-AOX | CO2-AOX + *pAOX1_ldhL + cyb2Δ + pMDH3_GYD1* | This study |
| CO2-LA-Mut- | CO2-LA-AOX + *aox2Δ::ADH2* | This study |
| CO2-LA-ADH | CO2-LA-Mut- + *Δaox2::ADH2 + pAOX1_ADH2* (multicopy clone – *ADH2* GCN=5) | This study |
| CO2-LA-ADH_Mit+Mxr | CO2-LA-ADH + *pFDH1_MIT1__pGAP_MXR1* (ENO locus) | This study |

### Construction of strains

The synthetic autotrophic *K. phaffii* strain described by Gassler et al. was used as a starting point for engineering Adh-based growth^10^. All plasmid were constructed by Golden Gate Assembly (GGA) as previously described^19^. When CRISPR-Cas9-mediated genome editing was employed, homologous donor DNA repair templates with 250-500 bp flanking homologous regions were provided, as previously described^18^. For the non-production strains, *AOX2* coding sequence was deleted through CRISPR-Cas9 using a homologous donor without the coding sequence (CO2-Mut-). A construct overexpressing *ADH2* either with or without a C-terminal PTS1 (SKL) tag was then integrated into the *RGI2* locus (CO2-ADH, CO2-ADH-Pts). Synthetic autotrophic *K. phaffii* strains described by Baumschabl et al. were used as starting points for engineering Adh-based production of lactate and itaconate^14^. The strain “cadA + mttA_medium_” was used directly and designated CO2-IA-AOX, and the strain “*ldhL* cyb2Δ” was altered by insertion of glyoxylate dehydrogenase (*GYD1)* from *Chlamydomonas reinhardtii* to avoid glycolate accumulation (CO2-LA-AOX). To produce Mut^-^ derivatives of both strains the coding sequence for *AOX2* was replaced with the native *ADH2* coding sequence (CO2-LA-Mut-, CO2-IA-Mut-).

A zeocin-selectable integration plasmid targeting the *AOX1*tt locus was constructed using the Golden PiCS modular cloning system and employed for multicopy integration of the pAOX_*ADH2* construct^19,41^. For selection of high copy number clones, 8 µg linearized donor DNA was used in the transformation, and transformants were plated on YPD containing 250, 500, or 1000 µg mL^−1^ zeocin. Colonies from the 500 and 1000 µg mL^−1^ plates were screened for *ADH2* gene copy number (GCN) and organic acid production, and the best performing strains were selected for further testing (CO2-IA-ADH, CO2-LA-ADH). To further improve lactic acid production in Adh-based strains, *MXR1* and *MIT1* described by Bachleitner et al. were assembled into a multi-gene plasmid targeting the *ENO1* locus, and inserted into CO2-LA-ADH using the standard transformation protocol described by Prielhofer et al.^19,32^.

### Gene copy number (GCN) determination

*ADH2* GCN was determined using quantitative PCR (qPCR). Reactions were prepared using 2× S’Green BlueMix (Biozyme Blue S’Green qPCR Kit), genomic DNA, nuclease-free water, and gene-specific primers according to the supplier’s instructions. All samples were analysed in technical quadruplets and no-template controls were included for each primer set. Quantitative PCR was performed on a Rotor-Gene real-time PCR system (Qiagen), and data were analysed using the comparative quantitation (CQ) method implemented in the Rotor-Gene software, with *ACT1* (PP7435_Chr3-0993) serving as the reference gene. Primer sequences used for *ADH2* were: forward 5′-CATGTCTCCAACTATCCCAACTACACAAAAG-3′ and reverse 5′-GACAGACACCGGAGTACTTAACG-3′. Primer sequences used for *ACT1* were: forward 5′-CCTGAGGCTTTGTTCCACCCATCT-3′ and reverse 5′-GGAACATAGTAGTACCACCGGACATAA-3′.

### Shake flask cultivations

Initial strain screenings were performed in 100 mL narrow-neck flasks inside a CO_2_ shaker-incubator where CO_2_ (5% or 10%) was supplied in addition to ambient air. A two-phase screening protocol was employed as described by Baumschabl et al.^14^. Briefly, biomass was generated in YPG precultures, after which an appropriate volume corresponding to a starting optical density (OD₆₀₀) of 4 was harvested, washed, and transferred to a new flask containing buffered YNB (3.4 g L^-1^, pH 6) supplemented with 10 g L^-1^ (NH_4_)_2_SO_4_ as the nitrogen source. Cultures were induced with 0.5% methanol (vol vol^-1^) at the start of the experiment and allowed to grow for 22-24 h before increasing the methanol concentration to either 1%, 2%, or 3% (vol vol^-1^). Every 1-3 days, the mass of each flask was measured and water was used to adjust for loss through evaporation before sampling. Optical density was measured from each sample, and the remaining volume was used for determination of methanol and organic acid concentrations by high-performance liquid chromatography (HPLC). Following analysis, methanol was added to adjust back to the target concentration.

### Kuhner TOM-shaker cultivations

Cultivations were performed in a shaker incubator using 300 mL wide-neck flasks (40–60 mL working volume) equipped with screw caps for controlled gas exchange. Caps were connected via gas lines to a Transfer Rate Online Measurement (TOM) module (Adolf Kühner AG, Birsfelden, Switzerland). TOM measurement cycles consisted of 30 min measurement followed by 120 min aeration. Ambient air with 5% CO_2_ was supplied at a rate of 11 mL min^-1^, and temperature set to either 25°C or 30°C. Oxygen transfer rate (OTR), CO₂ transfer rate (CTR), and respiratory quotient (RQ) were monitored and analyzed using the TOM-shaker software as previously described^29^. The same two-phase screening protocol used for shake-flask cultivations was applied, with offline measurements of optical density (OD₆₀₀), methanol, and organic acids. To determine dry cell weight (DCW) and establish the relationship between OD₆₀₀ and biomass concentration (g DCW L⁻¹), 2 mL or 10 mL samples were collected into pre-dried, pre-weighed Eppendorf tubes or volumetric flasks. Based on an OD₆₀₀-to-biomass conversion factor of 0.23 g CDW L⁻¹ per OD unit, specific CO₂ production (q_CO2_) and specific oxygen consumption (q_O2_) rates were calculated by normalizing CTR and OTR to the dry biomass estimated from OD₆₀₀ at each sampling point. At each sampling point, HPLC was used to quantify the concentrations of methanol, lactic and itaconic acid. Equations for calculation of further production parameters can be found in Table 4.

**Table 4.**
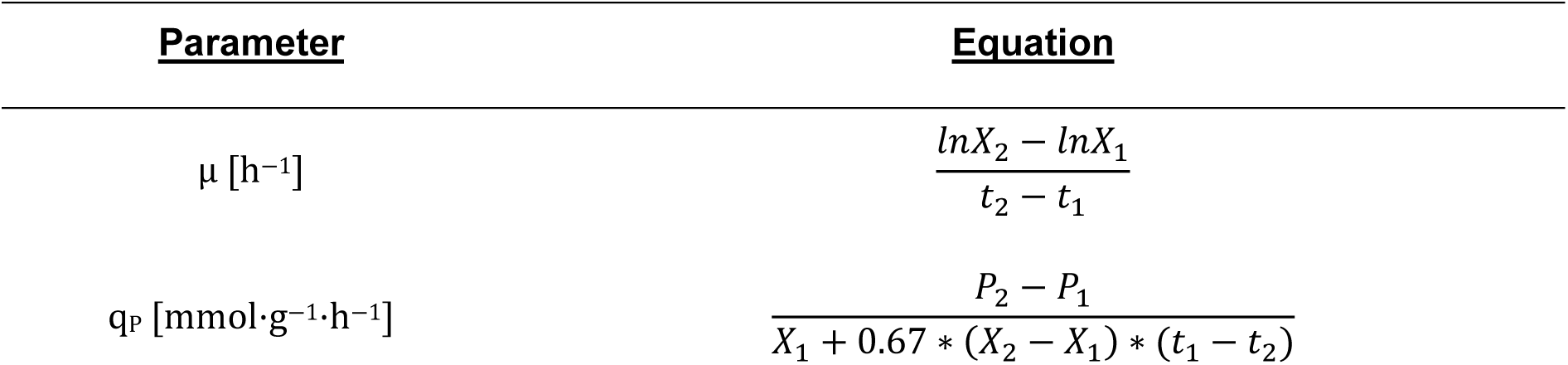
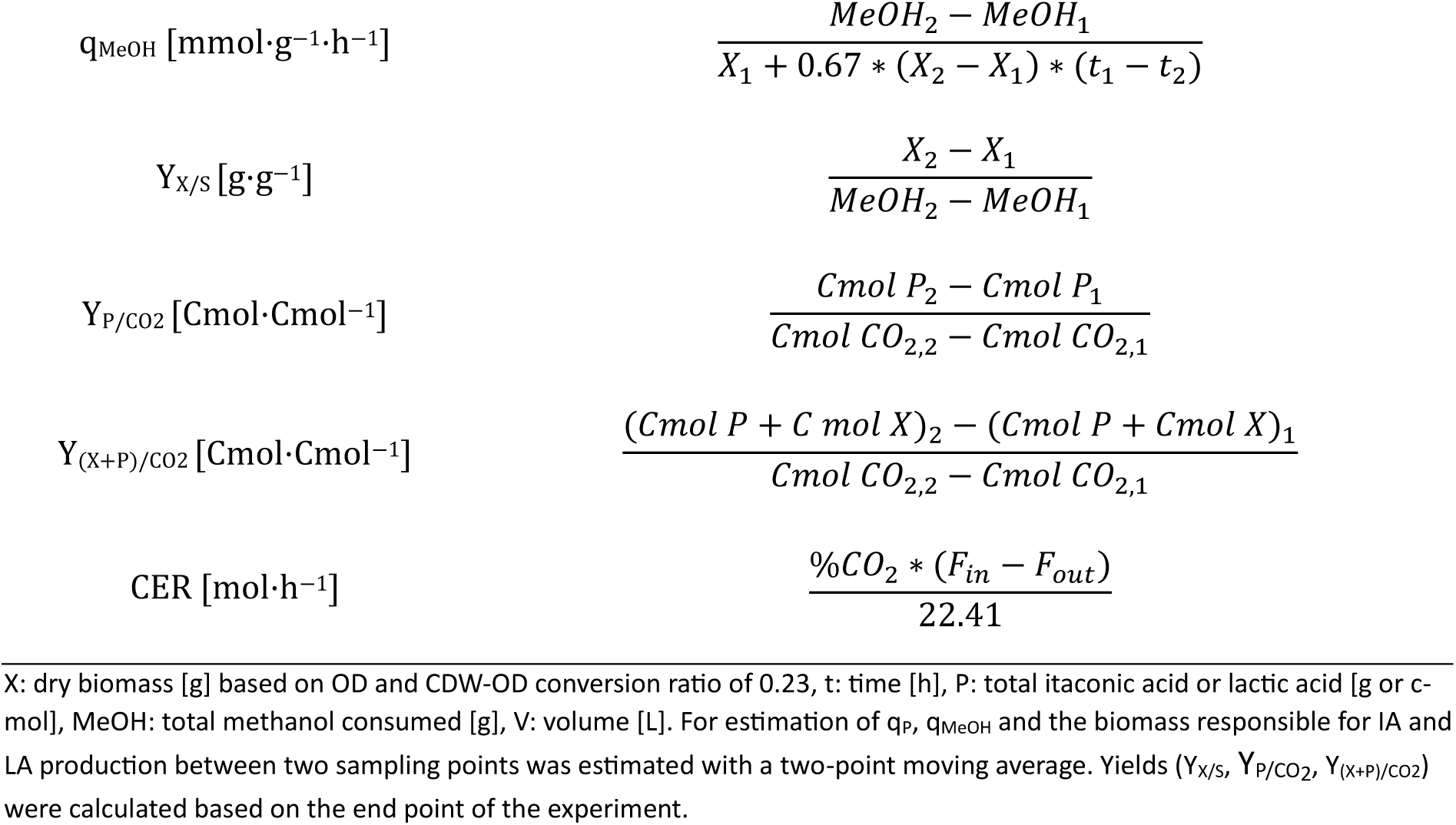
Equations used to calculate production parameters.

### Bioreactor Cultivations

Bioreactor cultivations were performed using 1.4 L DASGIP reactors (Eppendorf). Cultures were grown in YNB media supplemented with 10 g L^-1^ (NH_4_)_2_SO_4_ as the nitrogen source and buffered using 100 mmol L^-1^ phosphate buffer at pH 6 at 30°C. Reactors were inoculated from YPG precultures to an initial OD₆₀₀ of 4 in the presence of 0.5% (vol vol^-1^) methanol, and supplied with inlet gas containing 10% CO₂ and 10% O₂ at a flow rate of 6 sL h^-1^. Stirrer speed was set to 300 rpm and dissolved oxygen was not controlled. pH was controlled by automated addition of 2 mol L⁻¹ NaOH with a setpoint of 6.0. After the first sample (∼16 h) methanol concentration was adjusted to 1% (vol vol^-1^). Thereafter, samples were collected every 1–2 days for determination of OD₆₀₀, DCW, and methanol concentration. Following each sampling, methanol was replenished to maintain a target concentration of 1% (vol vol^-1^). Equations for calculation of CO_2_ evolution rate (CER) can be found in Table 4. To correct for baseline offsets in the off-gas analyser, for calculation of CER the measured CO₂,_out_ values were normalized by assuming an initial concentration of 5% prior to inoculation. To reduce high-frequency measurement noise, dissolved oxygen and CER trajectories were smoothed by index-based bin averaging (30 points per bin for dissolved oxygen and 1500 points per bin for CER), while optical density profiles were smoothed using locally estimated scatterplot smoothing (LOESS; span = 0.2) (Fig. 4).

### HPLC measurements

A Biorad Aminex HPX-87H HPLC column (300 × 7.8 mm) was used for the HPLC measurements^14^. H_2_SO_4_ at a concentration of 4mmol L^-1^ was used as mobile phase, with a 0.6 mL min^-1^ flow rate at 60°C. Itaconic acid was detected with a photodiode array detector (SPD-M20A, Shimadzu) at 254 nm. Lactic acid, and methanol concentrations were detected with a refraction index detector (RID-10A, Shimadzu). Samples were centrifuged, and the supernatant was mixed with H₂SO₄ (40 mmol L⁻¹ stock) to achieve a final concentration of 4 mmol L⁻¹. Following vortexing, samples were centrifuged at 16,100 × g for 5 min at room temperature and filtered through 0.22 μm filters into HPLC vials prior to analysis.

## Acknowledgements

The COMET center acib: Next Generation Bioproduction is funded by BMIMI, BMWET, SFG, Standortagentur Tirol, Government of Lower Austria and Vienna Business Agency in the framework of COMET – Competence Centers for Excellent Technologies. The COMET Funding Program is managed by the Austrian Research Promotion Agency FFG. We thank the Austrian Science Fund for support to D.M., C.M. and M.B. (Grant-DOI 10.55776/W1224, Doctoral Program on Biomolecular Technology of Proteins (BioToP)).

## Author contributions

D.M. and Ö.A. conceived and initiated the project. D.M., Ö.A. and C.M. designed the experiments. T.G. constructed the parental CO2-AOX strain and performed initial experiments with *ADH2* overexpression. C.M., M.B. and D.A. performed the further strain engineering. C.M. and L.L. carried out the shake flask experiments. C.M. and M.B. carried out the bioreactor experiment. C.M., Ö.A. and D.M. analysed the data. C.M., Ö.A. and D.M. wrote the manuscript. All authors read and approved the final manuscript.

