## Supplementary Information for "Alcohol dehydrogenase-mediated methanol dissimilation increases carbon efficiency in synthetic autotrophic yeast"

**Table S1.** Stoichiometric calculations for conversion of $CO_2$ to pyruvate in non-production strains

| Strain | Equations | MeOH /Cmol fixed | $CO_2$ net |
|---|---|---|---|
| | $3\ CO_2 + 7\ ATP + 5\ NADH \rightarrow PYR$ | | |
| CO2-AOX | $\underline{3.9\ MeOH \rightarrow 3.9\ CO_2 + 7.8\ NADH\ (= 19.5\ ATP)}$ | 1.30 | + 0.9 |
| | **$3.9\ MeOH \rightarrow PYR + 0.9\ CO_2$** | | |
| | $3\ CO_2 + 7\ ATP + 5\ NADH \rightarrow PYR$ | | |
| CO2-ADH | $\underline{2.6\ MeOH \rightarrow 2.6\ CO_2 + 7.8\ NADH\ (= 19.5\ ATP)}$ | 0.87 | - 0.4 |
| | **$2.6\ MeOH + 0.4\ CO_2 \rightarrow PYR$** | | |

ATP to NADH conversion factor of 2.5 was used assuming use of the malate-aspartate shuttle. PYR, pyruvate; MeOH, methanol.

**Table S2.** Cumulative production and consumption of carbon molecules in non-producing strains

| | | | **net production** | | **net consumption** |
|---|---|---|---|---|---|
| Strain | MeOH target [vol vol$^{-1}$] | temp. [°C] | X [mmol C] | $CO_2$ [mmol] | MeOH [mmol] |
| **CO2-Aox** | 1% | 30 | 2.62 | 9.87 | -13.4 |
| **CO2-Adh** | 1% | 30 | 2.36 | 4.31 | -7.59 |

$CO_2$ represents the net mol evolved, calculated from 'integral CTR' automatically calculated through the supplied software Kühner TOM shaker. A conversion factor of 0.23 was used for calculating DCW from OD. A conversion factor of 25.33 g Biomass per mol carbon was used for calculating carbon content in biomass.

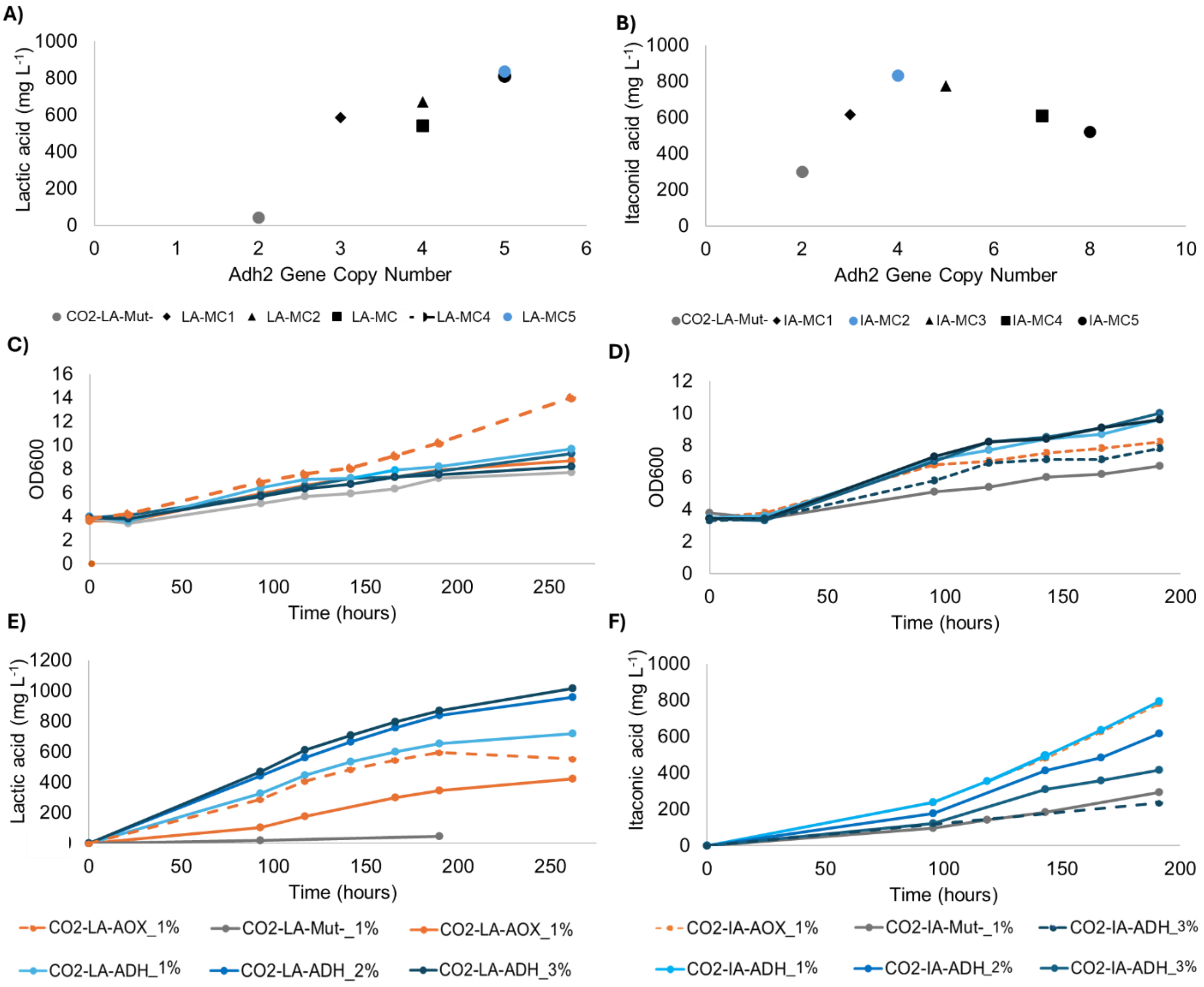


**Fig. S1| Improvement of organic acid production in Adh-based stains through metabolic engineering and process improvement**. (A,B) Cultivations performed in $CO_2$ shaker incubator at 30°C with 10% $CO_2$ and 2% methanol (vol vol$^{-1}$) for 200 h to identification of optimal *ADH2* gene copy number for organic acid production. The best performing strains indicated in blue were designated CO2-LA-ADH or CO2-IA-ADH and further tested for (C,D) growth and (E, F) organic acid production under different conditions. Cultivations performed at 25°C are indicated with a dotted line; cultivations performed at 30°C are indicated with a solid line. Methanol target (vol vol$^{-1}$) for the cultivation is indicated after the strain name. IA, itaconic acid; LA, lactic acid.

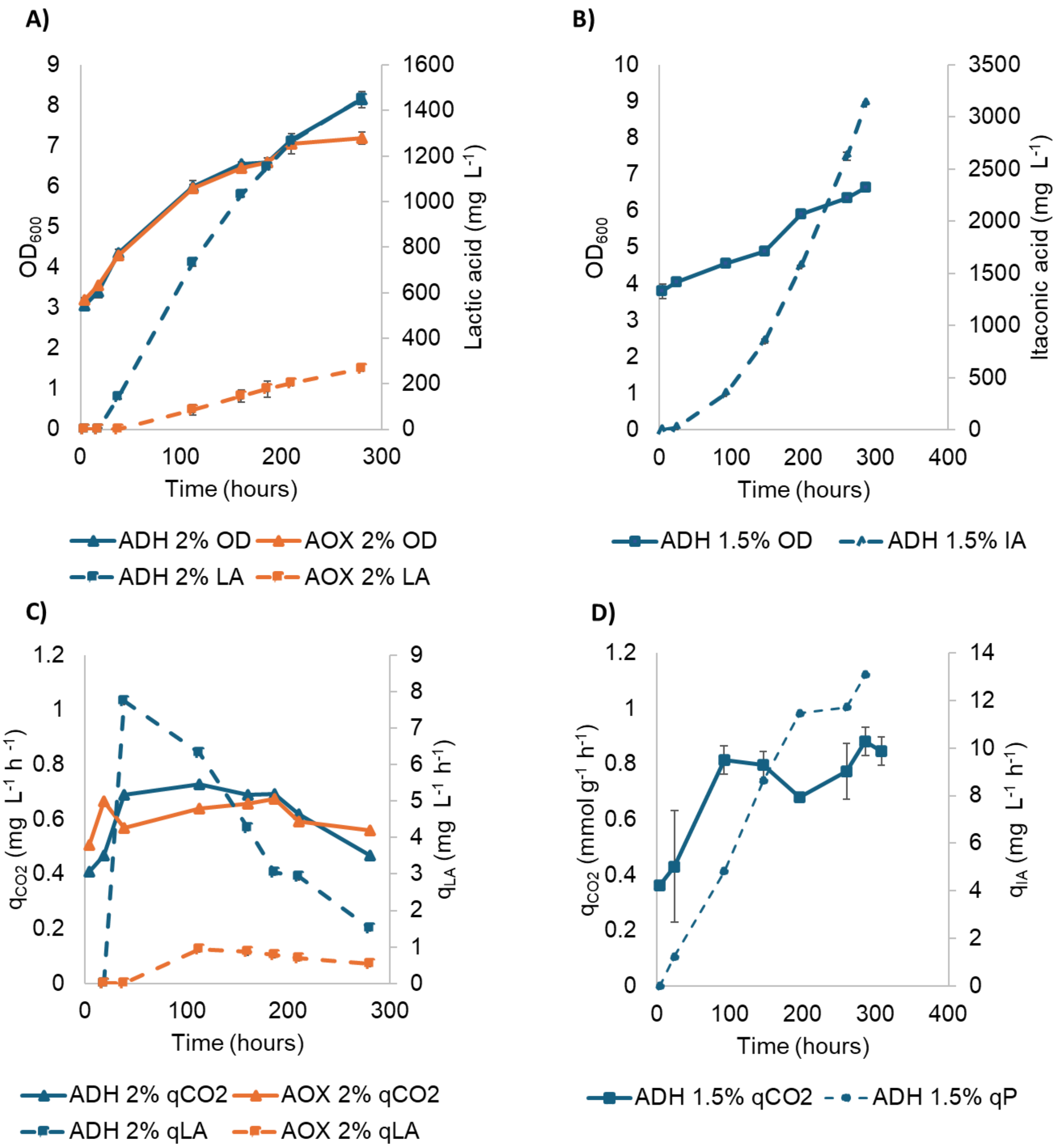


**Fig. S2| Comparison of growth, organic acid production, and net $CO_2$ evolution for production strains at elevated methanol concentrations in TOM-shaker unit.** Cultivation of CO2-LA-AOX and CO2-LA-ADH at 30°C with a 2% methanol (vol vol$^{-1}$) target with analysis of A) growth and lactic acid titer, and C) specific $CO_2$ production and lactic acid productivity. Cultivation of CO2-IA-ADH at 30°C with an increase in methanol target from 1% to 1.5% (vol vol$^{-1}$) at hour 182 h with analysis of B) growth and itaconic acid titer, and D) specific $CO_2$ production and itaconic acid productivity. performed to assess optical density and product formation for Aox-based producing strains at 30°C with 1% Methanol (vol vol$^{-1}$). IA, itaconic acid; LA, lactic acid. The mean of two individual cultivations for each strain is shown with standard deviation (±) is represented by or error bars.

**Table S3.** Stoichiometric calculations for conversion of $CO_2$ to organic acids in production strains

| Strain | Equations | MeOH /Cmol fixed | $CO_2$ net |
|---|---|---|---|
| CO2-IA-AOX | $4\ CO_2 + 8\ ATP + 5\ NADH \rightarrow OAA$<br>$3\ CO_2 + 7\ ATP + 4\ NADH \rightarrow PYR + CO_2$<br>$PYR + OAA \rightarrow ITA + CO_2$<br>$\underline{7.5\ MeOH \rightarrow 7.5\ CO_2 + 15\ NADH\ (= 37.5\ ATP)}$<br>**$7.5\ MeOH \rightarrow ITA + 2.5\ CO_2$** | 1.50 | + 2.5 |
| CO2-IA-ADH | $4\ CO_2 + 8\ ATP + 5\ NADH \rightarrow OAA$<br>$3\ CO_2 + 7\ ATP + 4\ NADH \rightarrow PYR + CO_2$<br>$PYR + OAA \rightarrow ITA + CO_2$<br>$\underline{5\ MeOH \rightarrow 5\ CO_2 + 15\ NADH\ (= 37.5\ ATP)}$<br>**$5\ MeOH \rightarrow ITA$** | 1.00 | 0.00 |
| CO2-LA-AOX | $3\ CO_2 + 7\ ATP + 6\ NADH \rightarrow LAC$<br>$\underline{4.4\ MeOH \rightarrow 4.4\ CO_2 + 8.8\ NADH\ (= 22\ ATP)}$<br>**$4.4\ MeOH \rightarrow LAC + 1.4\ CO_2$** | 1.47 | + 1.4 |
| CO2-LA-ADH | $3\ CO_2 + 7\ ATP + 6\ NADH \rightarrow PYR$<br>$\underline{2.93\ MeOH \rightarrow 2.93\ CO_2 + 8.8\ NADH\ (= 22\ ATP)}$<br>**$2.93\ MeOH + 0.07\ CO_2 \rightarrow LAC$** | 0.98 | - 0.07 |

ATP to NADH conversion factor was used assuming use of the malate-aspartate shuttle. PYR, pyruvate; MeOH, methanol; IA, itaconic acid; LA, lactic acid

**Table S4.** Cumulative production of carbon molecules in producing strains

| | | | net production | | |
|---|---|---|---|---|---|
| **Strain** | **MeOH target** | **temp.** | **X** | **$CO_2$** | **IA or LA** |
| | [vol vol$^{-1}$] | [°C] | [mmol C] | [mmol] | [mmol C] |
| **CO2-IA-AOX** | 1% | 30 | 0.50 | 12.7 | 3.17 |
| **CO2-IA-ADH** | 1% | 30 | 1.42 | 13.7 | 6.83 |
| **CO2-LA-AOX** | 1% | 30 | 2.01 | 9.35 | 0.48 |
| **CO2-LA-ADH** | 1% | 30 | 2.99 | 8.66 | 2.18 |

$CO_2$ represents the net mol evolved, calculated from 'integral CTR' automatically calculated through the supplied software Kühner TOM shaker. A conversion factor of 0.23 was used for calculating DCW from OD. A conversion factor of 25.33 g Biomass per mol carbon was used for calculating carbon content in biomass. IA, itaconic acid; LA, lactic acid.

**Table S5.** Average values of key parameters for Aox-based production strains at 25°C

| Strain | MeOH target | Temp | u | $q_P$ | $q_{CO2}$ | $q_{O2}$ |
|---|---|---|---|---|---|---|
| | [vol vol$^{-1}$] | [°C] | [$h^{-1}$] | [mg·$g^{-1}$·$h^{-1}$] | [mmol·$g^{-1}$·$h^{-1}$] | [mmol·$g^{-1}$·$h^{-1}$] |
| CO2-IA-AOX | 1% | 25 | 0.0103 | 2.95 | 0.706 | -1.38 |
| CO2-LA-AOX | 1% | 25 | 0.0104 | 0.585 | 0.815 | -1.48 |
| CO2-AOX | 1% | 25 | 0.0147 | -- | 0.802 | -1.69 |

$CO_2$ represents the net mol evolved, calculated from "integral CTR" automatically calculated through the supplied software Kühner TOM shaker. A conversion factor of 0.23 was used for calculating DCW from OD.

**Table S6.** Cumulative production of carbon molecules in Aox-based strains at 25°C

| | | | net production | | |
|---|---|---|---|---|---|
| **Strain** | **MeOH target** | **temp.** | **X** | **$CO_2$** | **IA or LA** |
| | [vol vol$^{-1}$] | [°C] | [mmol C] | [mmol] | [mmol C] |
| **CO2-AOX** | 1% | 25 | 7.56 | 24.2 | -- |
| **CO2-IA-AOX** | 1% | 25 | 6.44 | 28.24 | 4.39 |
| **CO2-LA-AOX** | 1% | 25 | 2.94 | 9.65 | 0.75 |

$CO_2$ represents the net mol evolved, calculated from 'integral CTR' automatically calculated through the supplied software Kühner TOM shaker. A conversion factor of 0.23 was used for calculating DCW from OD. A conversion factor of 25.33 g Biomass per mol carbon was used for calculating carbon content in biomass. IA, itaconic acid; LA, lactic acid

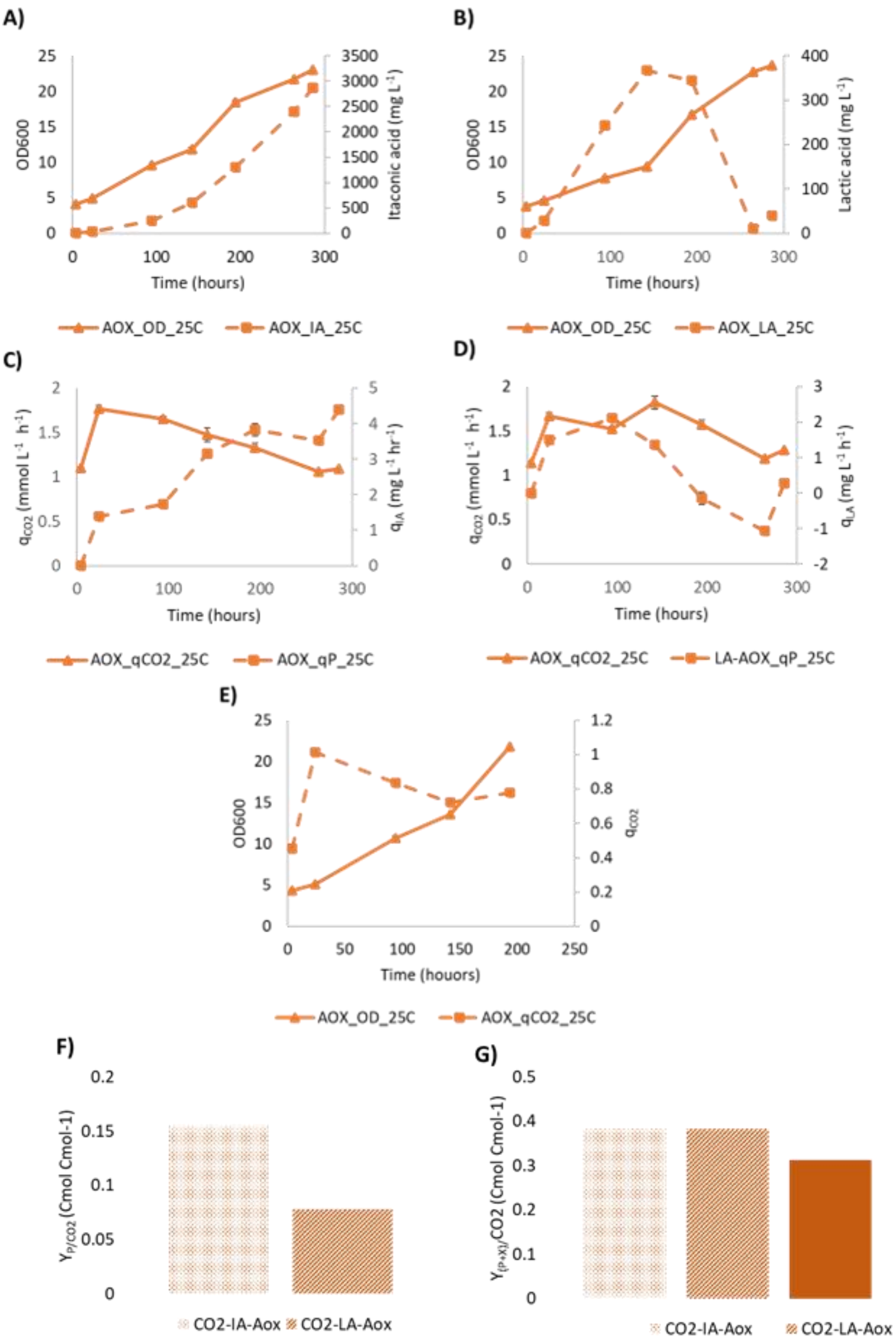

A)
OD600
Itaconic acid (mg L-1)
Time (hours)
AOX_OD_25C
AOX_IA_25C
B)
OD600
Lactic acid (mg L-1)
Time (hours)
AOX_OD_25C
AOX_LA_25C
C)
qCO2 (mmol L-1 h-1)
qIA (mg L-1 hr-1)
Time (hours)
AOX_qCO2_25C
AOX_qP_25C
D)
qCO2 (mmol L-1 h-1)
qLA (mg L-1 h-1)
Time (hours)
AOX_qCO2_25C
LA-AOX_qP_25C
E)
OD600
qCO2
Time (houors)
AOX_OD_25C
AOX_qCO2_25C
F)
YP/CO2 (Cmol Cmol-1)
CO2-IA-Aox
CO2-LA-Aox
G)
Y(P+X)/CO2 (Cmol Cmol-1)
CO2-IA-Aox
CO2-LA-Aox

**Fig. S3| Comparison of growth, organic acid production, and net $CO_2$ evolution for Aox-based strains in optimal conditions.** A TOM-shaker unit experiment performed to assess optical density and product formation for Aox-based producing strains at 25°C with 1% Methanol (vol $vol^{-1}$). A) growth and lactic acid titer, and C) specific $CO_2$ production and lactic acid productivity for CO2-LA-AOX strain. B) Growth and itaconic acid titer, and D) specific $CO_2$ production and itaconic acid productivity for CO2-IA-AOX strain. IA, itaconic acid; LA, lactic acid. The mean of two individual cultivations for each strain is shown with standard deviation (±) is represented by or error bars.

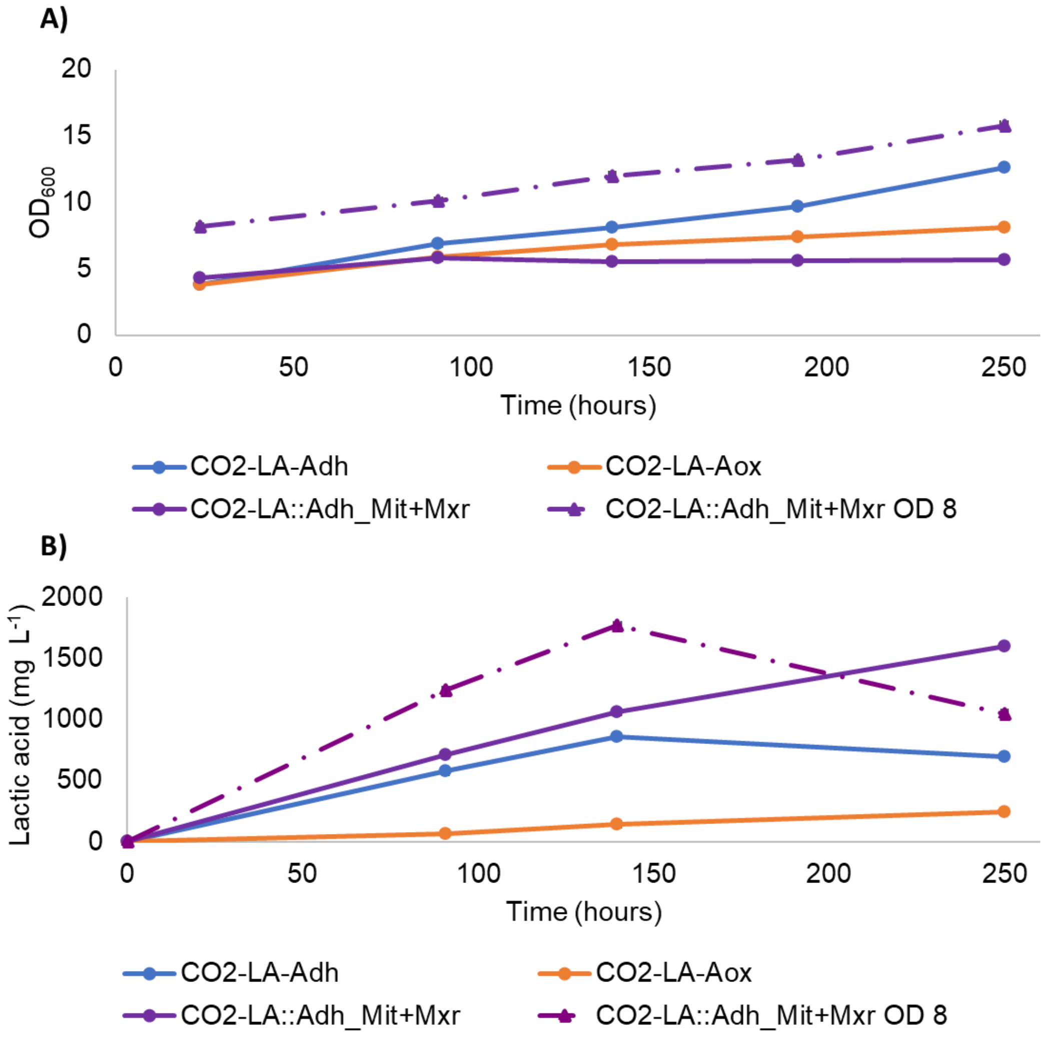


**Fig. S4|** Improvement of organic acid production in CO2-LA-ADH stains through overexpression of *MXR* and *MIT*. Cultivations were performed in a standard $CO_2$ shaker incubator with 10% $CO_2$ at 30°C. A) Optical density of strains and B) lactic acid concentrations. Parallel cultures for the same strain were tested with error bars representing standard deviation.
